# A modular, dynamic, DNA-based platform for regulating cargo distribution and transport between lipid domains

**DOI:** 10.1101/2021.03.02.433457

**Authors:** Roger Rubio-Sánchez, Simone Eizagirre Barker, Michal Walczak, Pietro Cicuta, Lorenzo Di Michele

## Abstract

Cell membranes regulate the distribution of biological machinery between phase-separated lipid domains to facilitate key processes including signalling and transport, which are among the life-like functionalities that bottom-up synthetic biology aims to replicate in artificial-cellular systems. Here, we introduce a modular approach to program partitioning of amphiphilic DNA nanostructures in co-existing lipid domains. Exploiting the tendency of different hydrophobic “anchors” to enrich different phases, we modulate the lateral distribution of our devices by rationally combining hydrophobes, and by changing nanostructure size and its topology. We demonstrate the functionality of our strategy with a bio-inspired DNA architecture, which dynamically undergoes ligand-induced reconfiguration to mediate cargo transport between domains *via* lateral re-distribution. Our findings pave the way to next-generation biomimetic platforms for sensing, transduction, and communication in synthetic cellular systems.

---

Biological membranes are highly heterogeneous, containing up to 20% of the protein content of a cell, and featuring hundreds of different lipid species.^1,2^ Such a degree of complexity evolved alongside the myriad of biological processes hosted and regulated by membranes, which include signalling, adhesion, trafficking, motility and division.^3^ Many of these functionalities rely on lateral co-localization of membrane proteins^4^ required for the emergence of signalling hubs,^5,6^ focal adhesions,^7^ and assemblies regulating membrane architecture to promote endo/exocytosis^8^ and cell division.^9^ Cells have evolved a variety of active and passive mechanisms to regulate the local composition of their membranes, overcoming the extreme molecular heterogeneity.^10–12^ Among these, proteo-lipid phase-separation is thought to play an important role in signalling and signal transduction. ^13^ In this process, nanoscale domains or *rafts* are believed to emerge, rich in sphingomyelins and sterols, and which are able to recruit or exclude membrane proteins based on their different affinities for raft and non-raft environments.^14,15^

Bottom-up synthetic biology aims at replicating functionalities typically associated to biological cells in microrobots designed *de novo*, or “artificial cells”.^16–18^ Just like their biological counterparts, many artificial-cell designs rely on semi-permeable membranes for their compartmentalization requirements, ^19–21^ which can be constructed from polymers^22^ and proteo-polymer systems,^23,24^ colloids^25,26^ and, more often, from synthetic lipid bilayers.^21^ However, with some remarkable exceptions^27–31^ reviewed in ref.,^32^ the membranes of artificial cells are often passive enclosures, lacking the complex functionalities of biological interfaces. A precise control over the local molecular makeup of synthetic lipid bilayers is therefore highly desirable, and a necessary stepping stone for the development of ever more sophisticated life-like responses in artificial cells.

DNA nanotechnology has demonstrated great potential as a means of creating responsive nanostructures that emulate biological architectures and functionalities, and are becoming increasingly popular constituents of artificial cellular systems.^33,34^ In many cases, biomimetic DNA nanostructures, rendered amphiphilic by hydrophobic tags, have been interfaced to synthetic lipid membranes to replicate the response of cell-membrane machinery. Examples include DNA architectures mediating artificial cell-adhesion and tissue formation,^35–41^ regulating transport,^42,43^ enabling signal transduction,^44^ and tailoring membrane curvature.^45,46^

Importantly, amphiphilic DNA nanostructures have been demonstrated to undergo preferential partitioning when anchored to phase-separated synthetic bilayers, an effect which is reminiscent of membrane-protein partitioning in rafts.^47–49^ The preference of DNA nanostructures for different lipid phases, and their degree of partitioning, have been shown to depend on the chemical identity of the hydrophobic anchors, membrane lipid composition, temperature, and solvent conditions.^40,50^ For instance, in ternary model membranes of DOPC/DPPC/cholesterol displaying coexistence of liquid ordered (*L*_o_) and liquid disordered (*L*_d_) phases,^1,51,52^ DNA constructs bearing a single cholesteryl-triethyleneglycol (TEG) anchor show a weak preference for *L*_o_, while duplexes end-functionalized with two cholesteryl-TEG anchors partition in *L*_o_ more prominently.^53,54^ Conversely, oligonucleotides functionalized with *α*-tocopherol preferentially localize within the *L*_d_ phases of ternary POPC/SM/cholesterol membranes.^55,56^ Moreover, dynamic control over the partitioning has also been exemplified, with DNA nanodevices that re-distribute between lipid phases following enzymatic cleavage^57^ or changes in ionic strength.^58^

The ability to influence partitioning of membrane-anchored DNA constructs makes them promising tools for engineering the molecular makeup and functionality of artificial cell membranes. Yet, we have effectively only started to scratch the surface of the massive design space of amphiphilic DNA nanostructures that, alongside the chemical identity of the hydrophobes, can be freely engineered with respect to their size, topology, and stimuli responsiveness.

Here we introduce a modular platform that fully exploits the design versatility of DNA nanotechnology to construct nanostructures whose ability to partition in the domains of phase-separated synthetic membranes can be precisely programmed. Besides enabling static membrane patterning, the nanodevices can be dynamically re-configured upon exposure to molecular cues, triggering re-distribution between domains and unlocking synthetic pathways for cargo transport, signalling and morphological adaptation in synthetic cellular systems.

Our nanostructures were constructed from synthetic DNA oligonucleotides, some of which were covalently linked to hydrophobic moieties to mediate anchoring to the membrane. Fluorescent tags (Alexa488, unless stated otherwise), were also included to monitor the distribution of the nanostructures *via* confocal microscopy. As depicted in Fig. 1 (left), we decorated the outer leaflet of phase-separated GUVs with the DNA constructs. Unless specified otherwise, GUVs were prepared from ternary lipid mixtures (DOPC/DPPC/cholestanol) to display *L*_d_-*L*_o_ coexistence. To enable fluorescence imaging, the vesicles were doped at 0.8% molar ratio with TexasRed-DHPE, which preferentially labels the *L*_d_ phase. At room temperature, the GUVs readily de-mixed into two macroscopic *L*_o_ and *L*_d_ domains. The resulting Janus-like morphology, shown in Fig. 1 (right), enabled the facile visualization and quantitation of the lateral distribution of the nanostructures.

**Figure 1:**
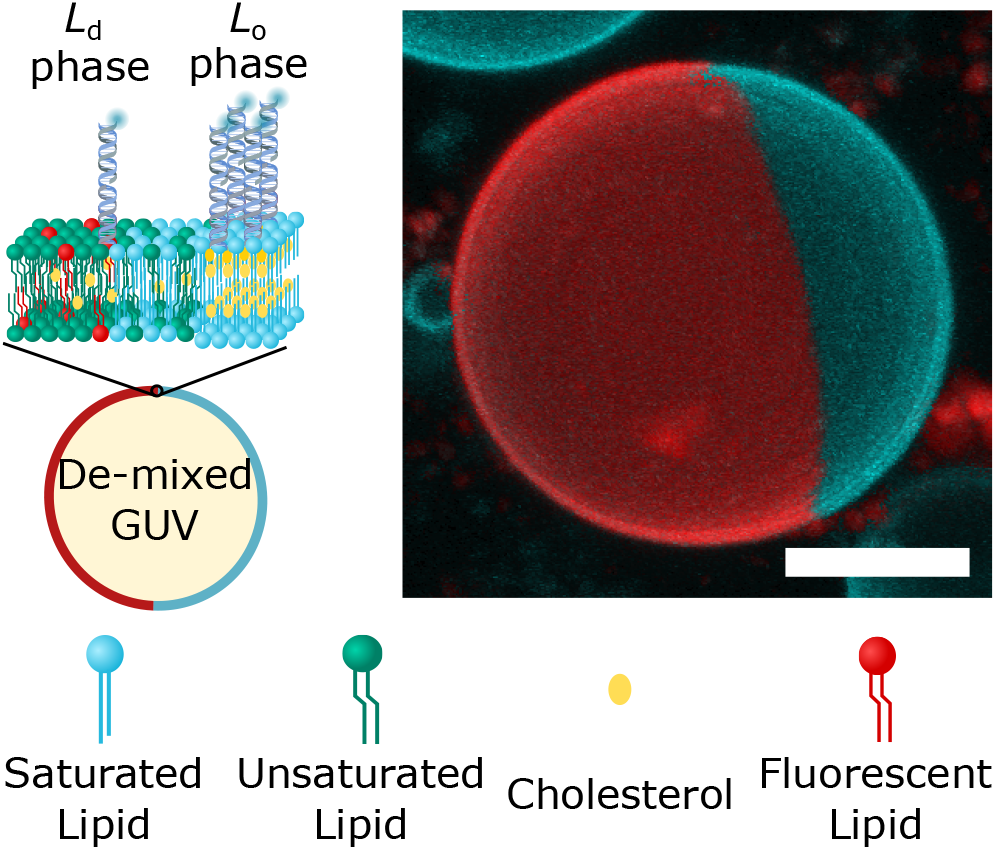
Membrane-anchored DNA nanostructures display preferential partitioning between the phases of demixed Giant Unilamellar Vesicles. Left: Schematic representation of a multi-component Giant Unilamellar Vesicle (GUV), prepared with saturated and unsaturated lipids mixed with sterols, displaying liquid-ordered (*L*_*o*_) and liquid-disordered (*L*_d_) phase co-existence. The outer leaflet of the membrane is functionalized with amphiphilic DNA nanostructures, exemplified here with constructs bearing two cholesterol anchors, which enrich their preferred (*L*_o_) phase. Right: 3D view of a de-mixed GUV from confocal z-stack as obtained from Volume viewer (FIJI^59^) using maximum projection. DNA nanostructures (cyan) partition to the *L*_o_ domain, while the *L*_d_ phase (red) is labelled with TexasRed-DHPE. Scale bar = 10 *μ*m.

To this end, equatorial confocal micrographs of the GUVs were collected and analyzed with a custom-built image processing pipeline, as described in detail in the SI (see Methods, and the associated Figs. S1 and S2), which determined the average fluorescence intensities of the constructs in the two phases: 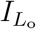 and 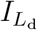. From these values, fractional intensities 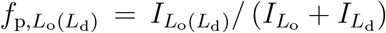 were extracted. Throughout this letter we refer to the fractional fluorescence intensity of nanostructures in the *L*_o_ phase, 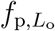, to gauge their partitioning tendency. Förster Resonance Energy transfer (FRET) between Alexa488 on the DNA and TexasRed on the lipids, alongside fluorescent-signal cross-talk, could in principle bias 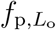 in certain partitioning states. To rule out this possibility, we performed dedicated experiments to determine the impact of both potential artefacts, as well as controls on GUVs that lack the TexasRed fluorophore, thus altogether eliminating the possible source of bias. Data in Fig. S3, and the associated Supplementary Discussion 1, confirm that FRET and fluorescence cross-talk carry a negligible impact on 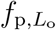.

Assuming that the recorded fluorescence intensities are proportional to nanostructure concentrations, a partitioning free energy can be calculated as 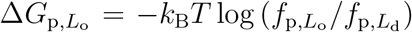. The latter is defined as the free energy change associated to relocating a single construct from the *L*_d_ to the *L*_o_ phase.

First, we applied our data-analysis pipeline on simple DNA duplexes featuring well characterized single cholesterol-TEG (sC) or single tocopherol (sT) anchors, as summarized in Fig. 2. Expectedly, while sC anchors led to a marginal *L*_o_-preference 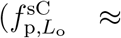 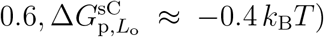, nanostructures featuring sT strongly partitioned in 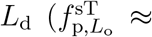 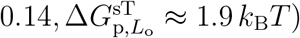.

**Figure 2:**
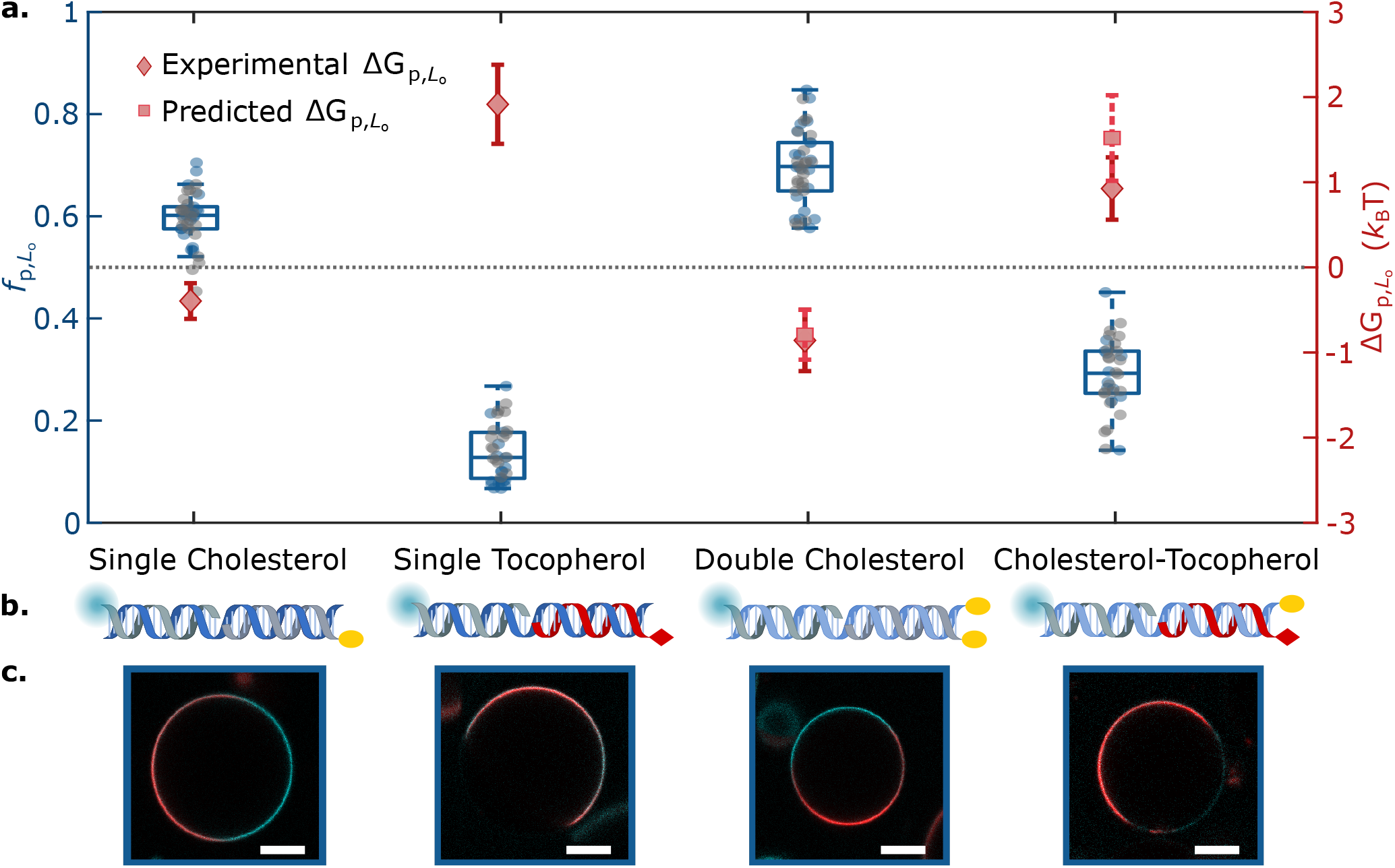
Lateral distribution of DNA nanostructures depends on anchor chemistry and combination. **a.** Fractional intensity 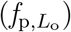 and free energy of partitioning 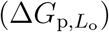 of DNA nanostructures in *L*_o_, conveyed by box-scatter plots and lozenges respectively, for two individual repeats (in blue/grey) of DNA-decorated GUV populations featuring different anchoring motifs: single cholesterol-TEG (sC), double cholesterol-TEG (dC), single tocopherol (sT), or a combination of single cholesterol-TEG and single tocopherol (sC+sT). For dC and sC+sT, squares indicate the predicted free energy values for combined anchors, determined from Eq. 1 using measured values of 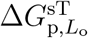 and 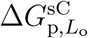. The dotted line indicates no partitioning 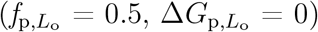. **b.** Schematic depictions of DNA duplexes bearing sC, sT, dC or sC+sT. **c.** Representative confocal micrographs of the DNA-functionalized GUVs. The *L*_d_ phase was labelled with TexasRed (red), and DNA constructs (cyan) with Alexa488. Scale bars = 10 *μ*m.

Importantly, as highlighted in Fig. S4, the size heterogeneity of electroformed GUVs does not affect the lateral distribution of DNA anchored species, as 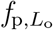 does not correlate with vesicle radius.

The substantial difference in the partitioning behaviors induced by sT and sC traces a route to program lateral distribution by combining multiple and/or different hydrophobic moieties within the same nanostructure. With this modular “mix and match” approach, we can indeed think of reinforcing partitioning in either lipid phase by increasing the number of cholesterol or tocopherol moieties on our nanodevices, or to access intermediate partitioning states with nanostructures featuring both moieties.

In the absence of (anti) co-operative effects and at sufficiently low construct concentrations, the partitioning free energy of a nanostructure featuring multiple hydrophobic moieties should be additive in the contributions from individual anchors

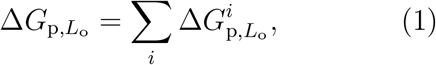

where the index *i* runs over all the anchors in the construct.

The simple relationship in Eq. 1 can guide the design of multi-anchor motifs and, in Fig. 2, we tested it on a duplex featuring two cholesterol-TEG anchors (dC). As expected, the dC motif displayed an enhanced preference for *L*_o_ domains compared to sC,^40,53^ with 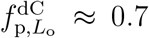. The measured partitioning free energy was 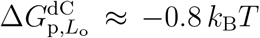, nearly identical to twice that of sC nanostructures. This quantitative agreement with the prediction of Eq. 1 suggests that, at least in this specific case, anchor co-operativity and other non-additive contributions negligibly affect partitioning.

When testing “chimeric” duplexes, bearing a tocopherol and a cholesterol anchor (sC+sT), we observed a pronounced preference for *L*_d_, consistent with the expectation that tocopherol should dominate in view of the stronger free energy shift associated with its partitioning (Fig. 2, 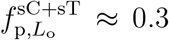). Here, the free energy prediction from Eq. 1 slightly overestimated the measured value of 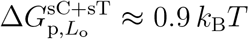 (statistically significant difference, *p* = 8.5 × 10^−11^, using the non-parametric Mann Whitney Wilconson Test). Since this non-additive behavior was not observed for the dC construct, we argue that it may result from the distinct chemical nature of the anchors in sC+dT and the consequent differences in their interactions with the surrounding lipids.

To further challenge our modular design approach, we studied the partitioning behavior of the constructs in Fig. 2 in a quaternary lipid mixture (DOPC/DPPC/cholestanol/cardiolipin). While this more complex mixture still displayed *L*_o_-*L*_d_ phase coexistence, the incorporation of the highly-unsaturated cardiolipin has been shown to enhance *L*_o_-partitioning, owing to the increased lateral stress in the *L*_d_ phase.^40^ The data, summarized in Fig. S5, largely confirmed the predictive power of Eq. 1, but some non-additive deviations are highlighted. Specifically, we observed that applying the rule in Eq. 1 led to an overestimation of the nanostructures’ tendency to localize within the *L*_o_ for both dC and sC+sT motifs. The difference in magnitude between non-additive free energy terms observed for ternary and quaternary lipid composition is only partially surprising, as one would expect (anti) co-operative effects to be highly sensitive to the lipid microenvironment of the anchors. For instance, one may speculate that the cardiolipin-rich *L*_d_ phase in the quaternary mixture may be better able to accommodate larger inclusions like those generated by two-anchor motifs (dC, sC+sT), which may in turn help relaxing the built-in lateral stress.^40^ This phenomenon may lead to a less pronounced *L*_o_ preference for two-anchor compared to single-anchor constructs.

Thanks to the design freedom of DNA nanotechnology, our modular strategy is not restricted to simple motifs with one or two anchors. We can indeed regard the duplex constructs in Fig. 2 as “anchoring modules”, and further combine them in larger nanostructures to expand the range of accessible partitioning states. For instance, as schematically depicted in Fig. 3, two anchoring modules were coupled by simply connecting them with a transversal, fluorescently labelled, linker duplex. For added conformational flexibility, 3-nt single-stranded (ss)DNA domains (*α* in Fig. 3) were included between the hydrophobically-modified and linker duplexes.

**Figure 3:**
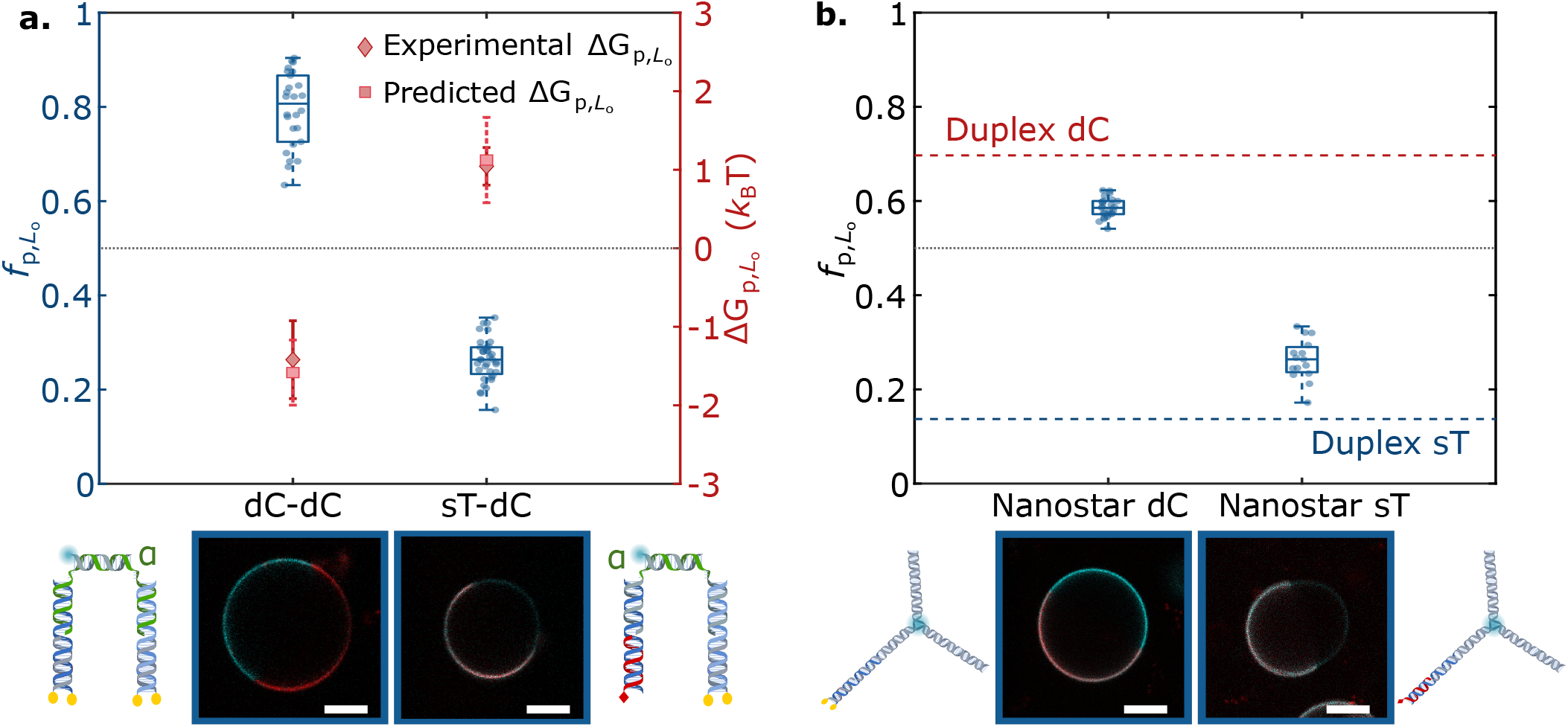
Anchor coupling and construct topology modulate DNA nanostructure partitioning. **a.** Top: Tunable lateral segregation of DNA architectures attained by coupling two sets of anchors: dC+dC and sT+dC. The partitioning behavior is demonstrated by the fractional intensity (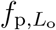, box-scatter plots) and free energy of partitioning (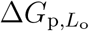, lozenges) in *L*_o_. Squares indicate the predicted free energy values for combined anchors, determined from Eq. 1 using measured values of 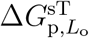 and 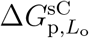 (Fig. 2). Bottom: Schematic representations of the nanostructures and representative confocal micrographs of decorated GUVs. **b.** Top: Effect of nanostructure size on partitioning, shown *via* 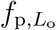 box-scatter plots of DNA nanostars bearing single tocopherol (sT) or double-cholesterol (dC) anchors, compared against the mean 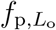 achieved by their duplex counterparts (dashed lines: red for dC; blue for sT). Bottom: Depiction of DNA nanostars alongside representative confocal micrographs of DNA-decorated GUVs. In both (**a** and **b**) plots, the dotted line indicates no partitioning 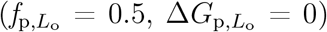. The *L*_d_ phase was labelled with Texas-Red (red), and constructs (cyan) with Alexa488 (DNA-DNA complexes) or fluorescein (nanostars). Scale bars = 10 *μ*m.

Two modular combinations were investigated: dC+dC and sT+dC. Notably, for both designs, the measured partitioning free energies could be quantitatively predicted by adding up the contributions of the individual anchor modules. The dC+dC construct, as expected, displays a very strong *L*_o_ preference, with 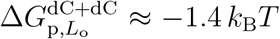 and 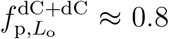. Similarly to the case of the sC+sT constructs, the partitioning of sT+dC nanostructures was dominated by the strong *L*_d_ preference of the tocopherol, with 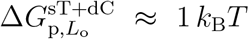 and 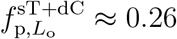.

Despite the remarkable accuracy of Eq. 1 in predicting partitioning states in multi-anchor constructs, we highlighted non-additive deviations. While some of these appear to correlate with the details of anchor and lipid chemistry (sC-sT in Fig. 2 and Fig. S5), and are thus difficult to control, other non-additive contributions can potentially be exploited to fine tune partitioning states around the baseline defined by anchor combination. For instance, the use of larger and more complex nanostructures may influence lateral segregation owing to steric or electrostatic interactions between the motifs. One may indeed expect that, for larger nanostructures, excluded volume effects may hinder accumulation in one specific phase, thus suppressing partitioning. Figure 3b summarizes the partitioning behavior of three-pointed DNA *nanostars* anchored to the bilayers using dC and sT modules. These motifs had roughly 4× the molecular weight of the smaller duplex architectures in Fig. 2 and, indeed, systematically displayed a reduced partitioning tendency. Moreover, Fig. S6 proves that the effect is not unique to the branched nanostructures, as linear duplexes anchored *via* dC also showed a general weakening in partitioning with increasing length. In further support to the hypothesis that steric nanostructure-nanostructure interactions may have an effect on partitioning, we performed measurements for smaller dC and sT duplexes over a wide range of (nominal) DNA-to-lipid molar ratios, and thus of surface densities of the motifs, as summarized in Fig. S7. We observed a near stationary 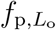 over a broad range of DNA/lipid ratios around the value used for all other experiments throughout this work (~ 4 × 10^−4^). For both dC and sT constructs, however, 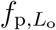 strongly deviated at high DNA/lipid ratios, approaching ≈ 0.5. Such a deviation hints at a saturation of the *L*_o_ and *L*_d_ phases, respectively, and the consequent impossibility for the nanostructures to further accumulate in those domains. Saturation occurred at higher DNA/lipid ratios for sT compared to dC, and we argue that this difference may arise from differences in the overall membrane affinity of the anchors. Indeed, while dC membrane insertion is effectively irreversible,^60^ anchoring *via* sT may be weaker, so that membrane-anchored sT duplexes coexist with a larger concentration of solubilized constructs, effectively reducing the surface density at fixed D NA/lipid ratio.

Our ability to program the domain partitioning of DNA constructs by linking different anchoring modules, as demonstrated in Fig. 3, can be combined with the dynamic reconfigurability of DNA nanostructures to reversibly trigger redistribution of a fluorescent cargo within the GUVs’ surfaces upon exposure to molecular cues. We demonstrate this effect with a nanostructure featuring both dC and sT anchoring modules, similar to that shown in Fig. 3 but in which the fluorescent dsDNA linker module (cargo) connecting the anchor duplexes can reversibly bind to or detach from either *via* toehold-mediated strand displacement^61^ – a mechanism that is reminiscent of two-component biological receptors undergoing ligand-induced dimerization.^5,6^ As sketched in Fig. 4a, we initiated our system from a configuration in which the fluorescent cargo was attached to the sT anchor, and thus localized in the *L*_d_ phase (State 1). Here, the fluorescent linker module was prevented from connecting to the dC anchoring module as its domain 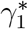, complementary to *γ*_1_ on the dC module, was protected by an Antifuel_1_ strand of domain sequence 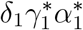. Note that the 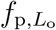 value recorded in this configuration matched exactly the one measured if the dC module was absent from the bilayer, marked by a dashed line in Fig. 4b, thus confirming the absence of binding to the dC module. Antifuel_1_ could be displaced upon addition of Fuel_1_, leading to the expo-sure of ^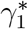^ on the linker module and its binding to the dC module. Upon acquiring this configuration, State 2, the nanostructures shifted the cargo towards *L*_o_. Addition of Antifuel_2_, with sequence 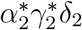, triggered the displacement of the linker from the sT module following a toeholding reaction initiated at domain *α*_2_, leading to the emergence of State 3 and a further cargo redistribution towards the *L*_o_ phase. Also in this configuration the 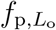 value aligned to that measured in the absence of sT anchors, indicating a near-complete progression of the toeholding reaction (dot-dash line in Fig. 4b). Finally, sequential addition of Fuel_2_ and Antifuel_1_ could reverse the systems’ configuration to States 2 and then 1. The last two steps pushed the fluorescent cargo back towards its initial *L*_d_ preference, but the 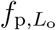 values remained slightly higher than those recorded at first. This incomplete reversibility may be due to partial inefficiencies in the reverse toeholding reactions, ^62^ or to a small thermodynamic unbalance that favors State 3 over State 1. Note that the system was allowed to fully equilibrate after the addition of fuel/antifuel strands prior to collecting the data in Fig. 4, as demonstrated by the measurements acquired at intermediate time-points and shown in Fig. S9a. In turn, Fig. S9b shows the time-evolution of 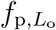 for an individual GUV, in which the nanostructures transition from State 1 to State 2 upon fuel addition. The equilibration timescales of ~5 minutes are likely limited by the diffusion of the fuel strand through the sample, given that, in order to prevent GUV drift and disruption, the fuel solution is gently added from the sample surface without any active mixing. The rates of toehold-mediated strand displacement and diffusion of the membrane-anchored constructs are expected to be comparatively faster (see discussion in the caption of Fig. S9). The redistribution recorded for our nanodevices is comparable with that of biological membrane-anchored agents involved *e.g.* in the recruitment of receptors upon T-cell activation^63^ or clathrin-mediated endocytosis,^8,64^ both of which range between tens to hundreds of seconds. Finally, note that States 1 and 3 displayed a lower tendency to partition in *L*_d_ and *L*_o_ (respectively) compared to their sT and dC duplex-analogues (Fig. 2). The shift is likely a consequence of the greater steric encumbrance of the responsive motifs, discussed above with respect to Fig. 3 and Fig. S6.

**Figure 4:**
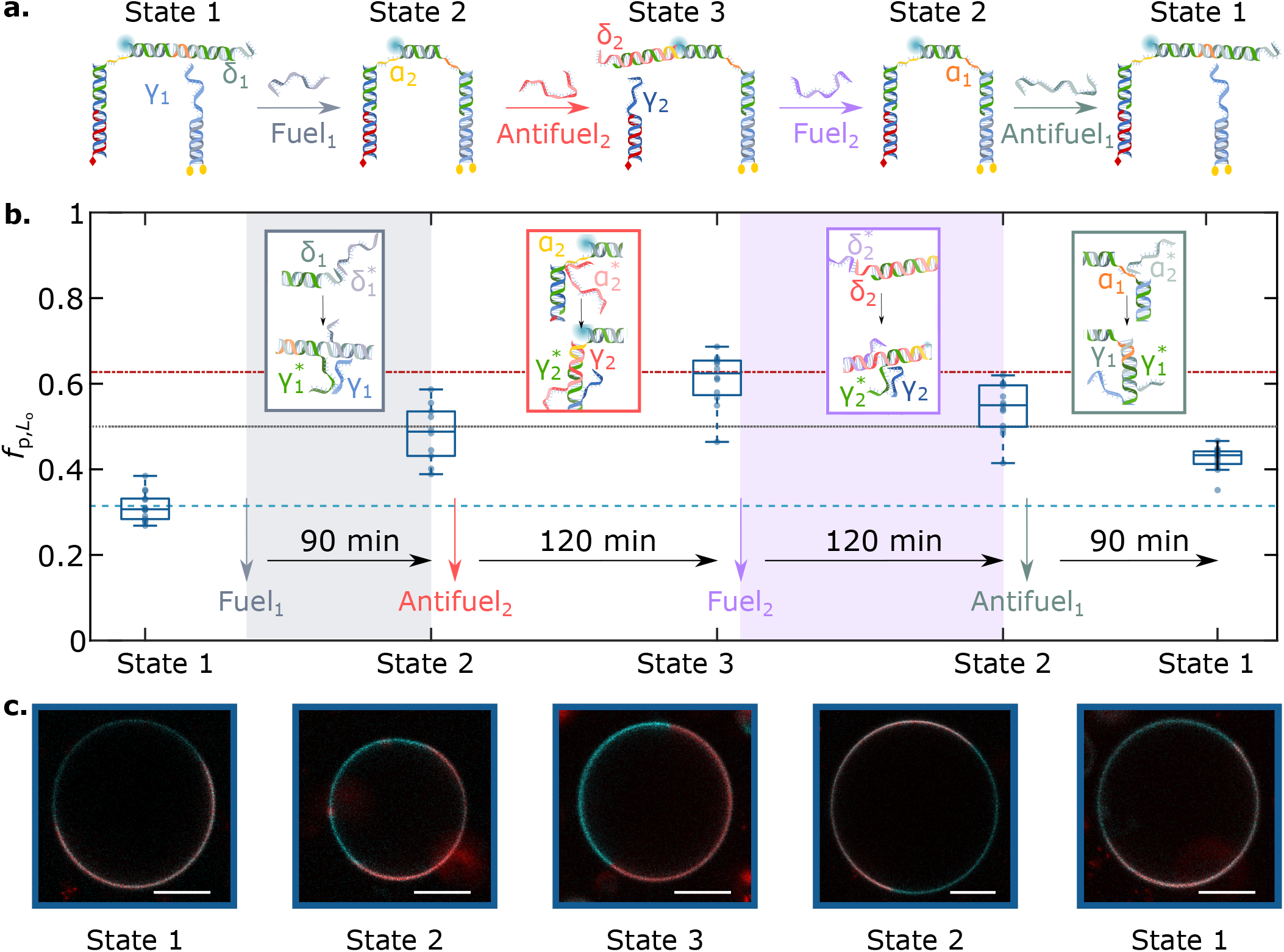
Toehold-mediated strand displacement enables cargo transport between lipid domains *via* lateral re-distribution to programmed partitioned states: **a.** Schematic depiction of the mechanism underpinning DNA nanostructure dimerization and cargo redistribution induced by the addition of Fuel/Antifuel strands, which exploit domains *δ*_1_*, α*_1_*, δ*_2_, and *α*_2_ as toe-holds. The correct response of the molecular circuit was tested with agarose gel electrophoresis, as summarized in Fig. S8. **b.** Evolution of the partitioning tendency of the fluorescent DNA element (panel **a**), conveyed by 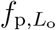 box-scatter plots, after a delay time for equilibration upon the addition of Fuel/Antifuel strands. The light blue dashed line marks the mean 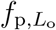 of the cargo hybridized to the dC-bearing module and in the absence of the sT-anchoring module, while the red dash-dot line that of the cargo hybridized to the sT-bearing module and in the absence of the dC anchoring module. The dotted line indicates no partitioning 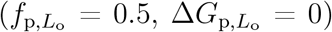. **c.** Representative confocal micrographs of DNA-functionalized GUVs over time, showcasing the distinct partitioned states attained by the system. The *L*_d_ phase was labelled with TexasRed (red), and DNA constructs (Alexa488) in cyan. Scale bars = 10 *μ*m.

In summary, we introduced a modular approach to engineer the lateral distribution of amphiphilic DNA nanostructures between co-existing phases of synthetic membranes. We exploited the ability of individual cholesterol and tocopherol anchors to induce partitioning in *L_o_* and *L*_d_ (respectively) and combined them to produce an array of multi-anchor nanodevices that span a broad range of partitioning behaviors from a ~ 80% preference for *L*_o_ to a ~ 85% partitioning in *L*_d_, as summarized in Fig. 5a. The comparison between measured partitioning free energies and those extracted from Eq. 1, shown in Fig. 5b, proves that to a good approximation 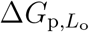 is additive in the contributions from individual anchors, thus offering a predictive design criterion. Small, non-additive effects contribute to a different extent depending on anchor combinations and lipid-membrane composition, hinting at (anti) co-operative effects dependent on system-specific chemistry, while excluded-volume effects emerged for bulkier motifs and higher nanostructure concentrations.

**Figure 5:**
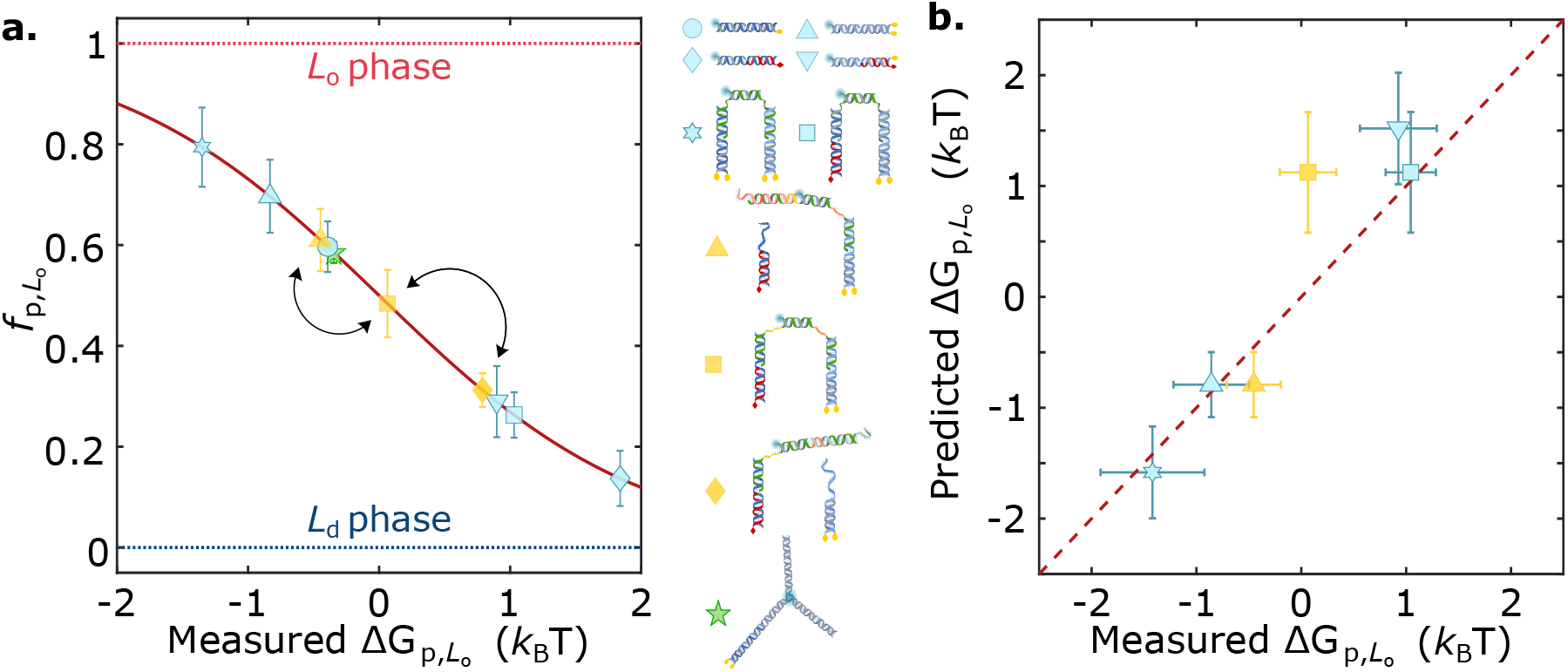
Static and dynamic engineering of partitioning states across the free-energy landscape by prescribing anchor combination and nanostructure morphology. **a.** Summary graph demonstrating the mean 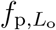 ± standard deviations of several membrane-associated DNA nanostructures as a function of their measured free energy of partitioning 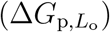. The effect of anchor coupling, size, and geometry is showcased. Arrows connect programmed partitioned states achieved by the responsive nanodevice discussed in Fig. 4. **b.** The partitioning free energy of DNA nanostructures is to a good approximation additive in contributions from individual anchoring motifs, as shown for several multi-anchor nanostructures with the means standard deviations of measured 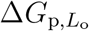 vs those predicted with Eq. 1.

The modularity of our design strategy enables dynamic control over the partitioning state by altering the anchor makeup of the nanostructures, as we showed with a proof-of-concept experiment in which fluorescent cargoes were reversibly transported across vesicle surfaces upon exposure to molecular cues. This strategy evokes that used by cells to control spatio-temporal localization of membrane proteins,^47–49,65^ and can be further extended to respond to other physico/chemical triggers by including stimuli-sensitive motifs such as aptamers,^66^ DNAzymes,^67^ G-quadruplex,^68^ pH-responsive motifs,^69^ and other functional DNA architectures.^70,71^

Our approach could be easily extended to include moieties other than cholesterol and tocopherol, such as alkyl chains,^40^ porphyrin,^72^ and azobenzene,^73^ which besides unlocking a broader range of partitioning states, may also enable responsiveness to a wider spectrum of stimuli. Similar design principles could even be applied to membrane-associated entities different from (amphiphilic) DNA nanostructures, such as peripheral and integral proteins, to program their co-localization in membrane domains.^74,75^

Our platform paves the way for the development of next-generation biomimetic DNA devices for the bottom-up engineering of life-like behaviors in synthetic cellular systems, which could in the long term find application in high-tech therapeutics and diagnostic solutions. For instance, the stimuli-triggered reshuffling of membrane-bound objects between lipid phases can enable highly sought behaviors such as signal transduction, communication and local membrane sculpting.^45^ Examples include stimuli-induced co-localization of *synthetic receptors* initiating artificial signalling cascades^44^ and herding of objects capable of influencing local membrane curvature, thus directing morphological re-structuring such as tubular growth^76^ and exosome/endosome budding.^46,77,78^

## Acknowledgement

R.R.S. acknowledges support from the Mexican National Council for Science and Technology (CONACYT, Grant No. 472427) and the Cambridge Trust. R.R.S and S.E.B also acknowledge funding from the EPSRC CDT in Nanoscience and Nanotechnology (NanoDTC, Grant No. EP/L015978/1 and EP/S022953/1, respectively). L.D.M. acknowledges funding from a Royal Society University Research Fellowship (UF160152) and from the European Research Council (ERC) under the Horizon 2020 Research and Innovation Programme (ERC-STG No 851667 NANOCELL). The authors thank B.M. Mognetti for discussions on the project, and D. Morzy for comments on the manuscript. All data in support of this work can be accessed free of charge at: https://doi.org/10.17863/CAM.64579.

## Supporting Information Available

Supporting Information contains the following items:

- Experimental Methods
- Supplementary Discussion
- Supplementary Figures
- DNA sequences of the nanostructures used throughout this work.

## References

(1) Heberle, F. A.; Feigenson, G. W. Phase Separation in Lipid Membranes. Cold Spring Harbor Perspectives in Biology 2011, 3, a004630.

(2) Lorent, J. H.; Levental, K. R.; Ganesan, L.; Rivera-Longsworth, G.; Sezgin, E.; Doktorova, M.; Lyman, E.; Levental, I. Plasma membranes are asymmetric in lipid unsaturation, packing and protein shape. Nature Chemical Biology 2020, 16, 644–652.

(3) Alberts, B. Molecular biology of the cell., sixth edit ed.; New York : Garland Science, 2017., 2017.

(4) Sezgin, E.; Levental, I.; Mayor, S.; Eggeling, C. The mystery of membrane organization: composition, regulation and roles of lipid rafts. Nature Reviews Molecular Cell Biology 2017, 18, 361–374.

(5) Akira, S.; Takeda, K. Toll-like receptor signalling. Nature Reviews Immunology 2004, 4, 499.

(6) Schlessinger, J. Receptor Tyrosine Kinases: Legacy of the First Two Decades. Cold Spring Harbor Perspectives in Biology 2014, 6, a008912.

(7) Acebrón, I. et al. Structural basis of Focal Adhesion Kinase activation on lipid membranes. The EMBO Journal 2020, 39, e104743.

(8) Haucke, V.; Kozlov, M. M. Membrane remodeling in clathrin-mediated endocytosis. Journal of Cell Science 2018, 131, jcs216812.

(9) Stoten, C. L.; Carlton, J. G. ESCRT-dependent control of membrane remodelling during cell division. Seminars in Cell & Developmental Biology 2018, 74, 50–65.

(10) Veatch, S. L.; Cicuta, P. In Critical Lipidomics: The Consequences of Lipid Miscibility in Biological Membranes BT - Physics of Biological Membranes; Bassereau, P., Sens, P., Eds.; Springer International Publishing: Cham, 2018; pp 141–168.

(11) McMahon, H. T.; Gallop, J. L. Membrane curvature and mechanisms of dynamic cell membrane remodelling. Nature 2005, 438, 590–596.

(12) Baumgart, T.; Capraro, B. R.; Zhu, C.; Das, S. L. Thermodynamics and Mechanics of Membrane Curvature Generation and Sensing by Proteins and Lipids. Annual Review of Physical Chemistry 2011, 62, 483–506.

(13) Pike, L. J. Lipid rafts : bringing order to chaos. Journal of Lipid Research 2003, 44, 655–667.

(14) Pralle, A.; Keller, P.; Florin, E.-L.; Simons, K.; Hörber, J. K. H. Sphin-golipid–Cholesterol Rafts Diffuse as Small Entities in the Plasma Membrane of Mammalian Cells. The Journal of Cell Biology 2000, 148, 997 LP – 1008.

(15) Lingwood, D.; Simons, K. Lipid Rafts As a Membrane-Organizing Principle. Science 2010, 327, 46 LP – 50.

(16) Buddingh’, B. C.; van Hest, J. C. M. Artificial Cells: Synthetic Compartments with Life-like Functionality and Adaptivity. Accounts of Chemical Research 2017, 50, 769–777.

(17) Noireaux, V.; Libchaber, A. A vesicle bioreactor as a step toward an artificial cell assembly. Proceedings of the National Academy of Sciences of the United States of America 2004, 101, 17669 LP – 17674.

(18) Noireaux, V.; Maeda, Y. T.; Libchaber, A. Development of an artificial cell, from self-organization to computation and self-reproduction. Proceedings of the National Academy of Sciences 2011, 108, 3473 LP – 3480.

(19) Elani, Y. Construction of membrane-bound artificial cells using microfluidics: a new frontier in bottom-up synthetic biology. Biochemical Society Transactions 2016, 44, 723 LP – 730.

(20) Karamdad, K.; Hindley, J. W.; Bolognesi, G.; Friddin, M. S.; Law, R. V.; Brooks, N. J.; Ces, O.; Elani, Y. Engineering thermoresponsive phase separated vesicles formed via emulsion phase transfer as a content-release platform. Chemical Science 2018, 9, 4851–4858.

(21) Trantidou, T.; Friddin, M.; Elani, Y.; Brooks, N. J.; Law, R. V.; Seddon, J. M.; Ces, O. Engineering Compartmentalized Biomimetic Micro- and Nanocontainers. ACS Nano 2017, 11, 6549–6565.

(22) Choi, H.-J.; Brooks, E.; Montemagno, C. D. Synthesis and characterization of nanoscale biomimetic polymer vesicles and polymer membranes for bio-electronic applications. Nanotechnology 2005, 16, S143–S149.

(23) Joesaar, A.; Yang, S.; Bögels, B.; van der Linden, A.; Pieters, P.; Kumar, B. V. V. S. P.; Dalchau, N.; Phillips, A.; Mann, S.; de Greef, T. F. A. DNA-based communication in populations of synthetic protocells. Nature Nanotechnology 2019, 14, 369–378.

(24) Ugrinic, M.; Zambrano, A.; Berger, S.; Mann, S.; Tang, T.-Y. D.; DeMello, A. Microfluidic formation of proteinosomes. Chem. Commun. 2018, 54, 287–290.

(25) Li, M.; Harbron, R. L.; Weaver, J. V. M.; Binks, B. P.; Mann, S. Electrostatically gated membrane permeability in inorganic protocells. Nature Chemistry 2013, 5, 529–536.

(26) Li, M.; Green, D. C.; Anderson, J. L. R.; Binks, B. P.; Mann, S. In vitro gene expression and enzyme catalysis in bio-inorganic protocells. Chem. Sci. 2011, 2, 1739–1745.

(27) Hindley, J. W.; Zheleva, D. G.; Elani, Y.; Charalambous, K.; Barter, L. M. C.; Booth, P. J.; Bevan, C. L.; Law, R. V.; Ces, O. Building a synthetic mechanosensitive signaling pathway in compartmentalized artificial cells. Proceedings of the National Academy of Sciences 2019, 116, 16711 LP – 16716.

(28) Altamura, E.; Milano, F.; Tangorra, R. R.; Trotta, M.; Omar, O. H.; Stano, P.; Mavelli, F. Highly oriented photosynthetic reaction centers generate a proton gradient in synthetic protocells. Proceedings of the National Academy of Sciences 2017, 114, 3837–3842.

(29) Litschel, T.; Ramm, B.; Maas, R.; Heymann, M.; Schwille, P. Beating Vesicles : Encapsulated Protein Oscillations Cause Dynamic Membrane Deformations. Angewandte Chemie International Edition 2018, 57, 16286–16290.

(30) Hindley, J. W.; Elani, Y.; McGilvery, C. M.; Ali, S.; Bevan, C. L.; Law, R. V.; Ces, O. Light-triggered enzymatic reactions in nested vesicle reactors. Nature Communications 2018, 9, 1093.

(31) Lee, K. Y.; Park, S.-J.; Lee, K. A.; Kim, S.-H.; Kim, H.; Meroz, Y.; Mahadevan, L.; Jung, K.-H.; Ahn, T. K.; Parker, K. K.; Shin, K. Photosynthetic artificial organelles sustain and control ATP-dependent reactions in a protocellular system. Nature Biotechnology 2018, 36, 530–535.

(32) Hindley, J. W.; Law, R. V.; Ces, O. Membrane functionalization in artificial cell engineering. SN Applied Sciences 2020, 2, 593.

(33) Seeman, N. C.; Sleiman, H. F. DNA nanotechnology. Nature Reviews Materials 2017, 3, 17068.

(34) Zhao, N.; Chen, Y.; Chen, G.; Xiao, Z. Artificial Cells Based on DNA Nanotechnology. ACS Applied Bio Materials 2020, 3, 3928–3934.

(35) Mognetti, B. M.; Cicuta, P.; Di Michele, L. Programmable interactions with biomimetic DNA linkers at fluid membranes and interfaces. Reports on Progress in Physics 2019, 82, 116601.

(36) Amjad, O. A.; Mognetti, B. M.; Cicuta, P.; Di Michele, L. Membrane Adhesion through Bridging by Multimeric Ligands. Langmuir 2017, 33, 1139–1146.

(37) Bachmann, S. J.; Kotar, J.; Parolini, L.; Šarić, A.; Cicuta, P.; Di Michele, L.; Mognetti, B. M. Melting transition in lipid vesicles functionalised by mobile DNA linkers. Soft Matter 2016, 12, 7804–7817.

(38) Parolini, L.; Mognetti, B. M.; Kotar, J.; Eiser, E.; Cicuta, P.; Di Michele, L. Volume and porosity thermal regulation in lipid mesophases by coupling mobile ligands to soft membranes. Nature Communications 2015, 6, 5948.

(39) Parolini, L.; Kotar, J.; Di Michele, L.; Mognetti, B. M. Controlling Self-Assembly Kinetics of DNA-Functionalized Liposomes Using Toehold Exchange Mechanism. ACS Nano 2016, 10, 2392–2398.

(40) Beales, P. A.; Nam, J.; Vanderlick, T. K. Specific adhesion between DNA-functionalized “Janus” vesicles: size-limited clusters. Soft Matter 2011, 7, 1747–1755.

(41) Beales, P. A.; Vanderlick, T. K. DNA as Membrane-Bound Ligand-Receptor Pairs: Duplex Stability Is Tuned by Intermembrane Forces. Biophysical Journal 2009, 96, 1554–1565.

(42) Burns, J. R.; Seifert, A.; Fertig, N.; Howorka, S. A biomimetic DNA-based channel for the ligand-controlled transport of charged molecular cargo across a biological membrane. Nature Nanotechnology 2016, 11, 152.

(43) Göpfrich, K.; Li, C.-Y.; Mames, I.; Bhamidimarri, S. P.; Ricci, M.; Yoo, J.; Mames, A.; Ohmann, A.; Winterhalter, M.; Stulz, E.; Aksimentiev, A.; Keyser, U. F. Ion Channels Made from a Single Membrane-Spanning DNA Duplex. Nano Letters 2016, 16, 4665–4669.

(44) Kaufhold, W. T.; Brady, R. A.; Tuffnell, J. M.; Cicuta, P.; Di Michele, L. Membrane Scaffolds Enhance the Responsiveness and Stability of DNA-Based Sensing Circuits. Bioconjugate Chemistry 2019, 30, 1850–1859.

(45) Franquelim, H. G.; Khmelinskaia, A.; Sobczak, J.-P.; Dietz, H.; Schwille, P. Membrane sculpting by curved DNA origami scaffolds. Nature Communications 2018, 9, 811.

(46) Journot, C. M. A.; Ramakrishna, V.; Wallace, M. I.; Turberfield, A. J. Modifying Membrane Morphology and Interactions with DNA Origami Clathrin-Mimic Networks. ACS Nano 2019, 13, 9973–9979.

(47) Wang, T.-Y.; Leventis, R.; Silvius, J. R. Partitioning of Lipidated Peptide Sequences into Liquid-Ordered Lipid Domains in Model and Biological Membranes. Biochemistry 2001, 40, 13031–13040.

(48) Weise, K.; Triola, G.; Janosch, S.; Waldmann, H.; Winter, R. Visualizing association of lipidated signaling proteins in heterogeneous membranes Partitioning into subdomains, lipid sorting, interfacial adsorption, and protein association. Biochimica et Biophysica Acta (BBA) - Biomembranes 2010, 1798, 1409–1417.

(49) Weise, K.; Kapoor, S.; Denter, C.; Nikolaus, J.; Opitz, N.; Koch, S.; Triola, G.; Herrmann, A.; Waldmann, H.; Winter, R. Membrane-Mediated Induction and Sorting of K-Ras Microdomain Signaling Platforms. Journal of the American Chemical Society 2011, 133, 880–887.

(50) Beales, P. A.; Vanderlick, T. K. Application of nucleic acid–lipid conjugates for the programmable organisation of liposomal modules. Advances in Colloid and Interface Science 2014, 207, 290–305.

(51) Veatch, S. L.; Keller, S. L. Separation of Liquid Phases in Giant Vesicles of Ternary Mixtures of Phospholipids and Cholesterol. Biophysical Journal 2003, 85, 3074–3083.

(52) Veatch, S. L.; Keller, S. L. Miscibility Phase Diagrams of Giant Vesicles Containing Sphingomyelin. Phys. Rev. Lett. 2005, 94, 148101.

(53) Beales, P. A.; Vanderlick, T. K. Partitioning of Membrane-Anchored DNA between Coexisting Lipid Phases. The Journal of Physical Chemistry B 2009, 113, 13678–13686.

(54) Talbot, E. L.; Parolini, L.; Kotar, J.; Di Michele, L.; Cicuta, P. Thermal-driven domain and cargo transport in lipid membranes. Proceedings of the National Academy of Sciences 2017, 114, 846 LP – 851.

(55) Bunge, A.; Kurz, A.; Windeck, A.-K.; Korte, T.; Flasche, W.; Liebscher, J.; Herrmann, A.; Huster, D. Lipophilic Oligonucleotides Spontaneously Insert into Lipid Membranes, Bind Complementary DNA Strands, and Sequester into Lipid-Disordered Domains. Langmuir 2007, 23, 4455–4464.

(56) Kurz, A.; Bunge, A.; Windeck, A.-K.; Rost, M.; Flasche, W.; Arbuzova, A.; Strohbach, D.; Müller, S.; Liebscher, J.; Huster, D.; Herrmann, A. Lipid-Anchored Oligonucleotides for Stable Double-Helix Formation in Distinct Membrane Domains. Angewandte Chemie International Edition 2006, 45, 4440–4444.

(57) Schade, M.; Knoll, A.; Vogel, A.; Seitz, O.; Liebscher, J.; Huster, D.; Herrmann, A.; Arbuzova, A. Remote Control of Lipophilic Nucleic Acids Domain Partitioning by DNA Hybridization and Enzymatic Cleavage. Journal of the American Chemical Society 2012, 134, 20490–20497.

(58) Czogalla, A.; Petrov, E. P.; Kauert, D. J.; Uzunova, V.; Zhang, Y.; Seidel, R.; Schwille, P. Switchable domain partitioning and diffusion of DNA origami rods on membranes. Faraday Discussions 2013, 161, 31–43.

(59) Schindelin, J. et al. Fiji: an open-source platform for biological-image analysis. Nature Methods 2012, 9, 676–682.

(60) Pfeiffer, I.; Höök, F. Bivalent Cholesterol-Based Coupling of Oligonucletides to Lipid Membrane Assemblies. Journal of the American Chemical Society 2004, 126, 10224–10225.

(61) Zhang, D. Y.; Winfree, E. Control of DNA Strand Displacement Kinetics Using Toehold Exchange. Journal of the American Chemical Society 2009, 131, 17303–17314.

(62) Chen, R. P.; Blackstock, D.; Sun, Q.; Chen, W. Dynamic protein assembly by programmable DNA strand displacement. Nature Chemistry 2018, 10, 474–481.

(63) Bini, L.; Pacini, S.; Liberatori, S.; Valensin, S.; Pellegrini, M.; RaggiaschiI, R.; Pallani, V.; Baldari, C. T. Extensive temporally regulated reorganization of the lipid raft proteome following T-cell antigen receptor triggering. Biochemical Journal 2003, 369, 301–309.

(64) Taylor, M. J.; Perrais, D.; Merrifield, C. J. A High Precision Survey of the Molecular Dynamics of Mammalian Clathrin-Mediated Endocytosis. PLOS Biology 2011, 9, e1000604.

(65) Rocks, O.; Peyker, A.; Bastiaens, P. I. H. Spatio-temporal segregation of Ras signals: one ship, three anchors, many harbors. Current Opinion in Cell Biology 2006, 18, 351–357.

(66) Del Grosso, E.; Ragazzon, G.; Prins, L. J.; Ricci, F. Back Cover : Fuel-Responsive Allosteric DNA-Based Aptamers for the Transient Release of ATP and Cocaine (Angew. Chem. Int. Ed. 17/2019). Angewandte Chemie International Edition 2019, 58, 5772.

(67) Lu, Y.; Liu, J. Functional DNA nanotechnology: emerging applications of DNAzymes and aptamers. Current Opinion in Biotechnology 2006, 17, 580–588.

(68) Cozzoli, L.; Gjonaj, L.; Stuart, M. C. A.; Poolman, B.; Roelfes, G. Responsive DNA G-quadruplex micelles. Chem. Commun. 2018, 54, 260–263.

(69) Idili, A.; Vallée-Bélisle, A.; Ricci, F. Programmable pH-Triggered DNA Nanoswitches. Journal of the American Chemical Society 2014, 136, 5836–5839.

(70) Ma, L.; Liu, J. Catalytic Nucleic Acids: Biochemistry, Chemical Biology, Biosensors, and Nanotechnology. iScience 2020, 23, 100815.

(71) Le Vay, K.; Salibi, E.; Song, E. Y.; Mutschler, H. Nucleic Acid Catalysis under Potential Prebiotic Conditions. Chemistry – An Asian Journal 2019, 15, 214–230.

(72) Burns, J. R.; Göpfrich, K.; Wood, J. W.; Thacker, V. V.; Stulz, E.; Keyser, U. F.; Howorka, S. Lipid-Bilayer-Spanning DNA Nanopores with a Bifunctional Porphyrin Anchor. Angewandte Chemie International Edition 2013, 52, 12069–12072.

(73) Hernández-Ainsa, S.; Ricci, M.; Hilton, L.; Aviñó, A.; Eritja, R.; Keyser, U. F. Controlling the Reversible Assembly of Liposomes through a Multistimuli Responsive Anchored DNA. Nano Letters 2016, 16, 4462–4466.

(74) Lorent, J. H.; Diaz-Rohrer, B.; Lin, X.; Spring, K.; Gorfe, A. A.; Levental, K. R.; Levental, I. Structural determinants and functional consequences of protein affinity for membrane rafts. Nature Communications 2017, 8, 1219.

(75) Lin, X.; Gorfe, A. A.; Levental, I. Protein Partitioning into Ordered Membrane Domains: Insights from Simulations. Biophysical journal 2018, 114, 1936–1944.

(76) Talbot, E. L.; Kotar, J.; Di Michele, L.; Cicuta, P. Directed tubule growth from giant unilamellar vesicles in a thermal gradient. Soft Matter 2019, 15, 1676–1683.

(77) Booth, A.; Marklew, C. J.; Ciani, B.; Beales, P. A. In Vitro Membrane Remodeling by ESCRT is Regulated by Negative Feedback from Membrane Tension. iScience 2019, 15, 173–184.

(78) Booth, A.; Marklew, C.; Ciani, B.; Beales, P. A. The influence of phosphatidylserine localisation and lipid phase on membrane remodelling by the ESCRT-II/ESCRT-III complex. Faraday Discussions 2020,

