## Supporting Information for "A modular, dynamic, DNA-based platform for regulating cargo distribution and transport between lipid domains"

#### Methods

**Multicomponent Phase-separating Giant Unilamellar Vesicles:** Vesicles were prepared *via* electroformation[1–3] in a 300 mM sucrose solution in MilliQ water, using 1,2-dioleoyl-sn-glycero-3-phosphocholine [DOPC] (Avanti Polar Lipids; chain melting temperature -17°C), 1,2-dipalmitoyl-sn-glycero-3-phosphocholine [DPPC] (Avanti Polar Lipids; chain melting temperature 41°C), cholestanol (Sigma-Aldrich) and cardiolipin (Sigma-Aldrich) in relevant molar ratios. The composition of ternary GUVs used in the main text was DOPC:DPPC:Chol = 4:4:2, while that of quaternary GUVs (Fig. S5) was DOPC:DPPC:Chol:Cardiolipin

= 2.75:3.75:2.5:1. Lipid mixtures were doped at 0.8% molar ratio with fluorescent lipid TexasRed-DHPE (Invitrogen), which preferentially co-localizes in the $L_\mathrm{d}$ phase. Briefly, indium tin oxide (ITO) slides (Visiontek Systems Ltd) were cleaned by sonication cycles of isopropanol/milli Q water for 15 min and dried under a gentle nitrogen flow. The slides were subsequently heated up to 60°C, and 45 $\mu$L of lipid mixtures (4 mg$\cdot$ mL$^{-1}$) were spread on the conducting side using a clean glass coverslip. The slides, after resting for 1 hour under vacuum in a dry silica dessicator, were assembled into an electroformation chamber by coupling them with a $\sim$ 1 mm thick polydimethylsiloxane (PDMS) spacer enclosing approximately 300 $\mu$L of degassed sucrose buffer. The chambers were placed in a pre-heated oven at $T > 60$°C, coupled to a frequency generator with clamps and subjected to a sinusoidal alternating current (AC) with voltage amplitude of 2V. Initially, the frequency was set to 10 Hz for a period of 2 hrs and then to 2 Hz for 1 hr. Vesicles were then retrieved and stored at room temperature in the dark to prevent photo-bleaching and photo-oxidation.

**Quantification of Lipid Concentration in Electroformed Vesicles:** Electroformed GUVs were subjected to a phosphatidylcholine (PC) enzymatic assay (Sigma Aaldrich), as per the manufacturer's technical bulletin, to determine lipid concentration and compute the DNA/lipid molar ratio used *e.g.* in Fig. S7. PC concentration was determined *via* colorimetric measurements by acquiring absorbance at 570 nm ($\mathrm{A}_{\lambda=570\mathrm{nm}}$) with a FLUOstar Omega plate reader. Samples were diluted in a reaction mix containing assay buffer, PC hydrolysis enzyme, a development mix, and fluorescent peroxidase substrate as indicated by the manufacturer. After incubation for $\sim$30 min, $\mathrm{A}_{\lambda=570\mathrm{nm}}$ was measured for an hour, and averaged for subsequent calculations. All samples were blanked against absorbance measurements of vesicles in a reaction mix lacking the enzyme. PC concentration was determined from a calibration curve run on standard samples of PC as instructed by the manufacturer.

**DNA Nanostructures Assembly and Manipulation:** DNA constructs were designed using a nucleic acid design and analysis software (NUPACK[4]) based on previously reported modules.[5] DNA oligonucleotides were purchased lyophilized (Integrated DNA Technologies [IDT], Eurogentec, and Biomers), purified by the supplier with high-performance liquid chromatography (HPLC), and resuspended in Tris-Ethylenediaminetetraacetic acid (EDTA) buffer (1× TE: 10 mM Tris + 1 mM EDTA, pH 8.0) to a final concentration of 100 $\mu$M. DNA constructs self-assembled with a slow quenching temperature ramp on a TC-512 thermal cycler (95°C down to 4°C at a rate of -0.5°C min $^{-1}$) in a buffer containing 1× TE + 100 mM NaCl.

**Vesicle Functionalization with DNA nanostructures:** DNA constructs (16.7 $\mu$L) were mixed with 9.2 $\mu$L of GUVs in 57.4 $\mu$L of a correcting buffer to result in an iso-osmolar mixture containing 1× TE + 100 mM NaCl + 87 mM Glucose to a final DNA/lipid molar ratio of $\sim 4 \times 10^{-4}$ (unless stated otherwise). The mixture was left under rotation overnight and subsequently stored at room temperature in the dark to avoid photobleaching. In the case of measurements on the responsive nanodevice, fuel/antifuel strands were added sequentially in iso-osmolar buffers (as above) at 5, 10, 15, and 20× excess with respect to the anchoring modules and the fluorescent cargo.

**Agarose Gel Electrophoresis (AGE):** Gels were prepared at 2% (weight) of agarose in Tris-Borate-EDTA (TBE, Sigma Aldrich - 89 mM Tris-borate, 2 mM EDTA, pH 8.3) buffer. The mixture was dissolved *via* heating. SYBR Safe DNA gel stain (Invitrogen) was added at 0.1% (volume) and gently dissolved through swirling. The mixture was casted to a thickness of approximately 5 mm and allowed to set for 1 hour. Subsequently, the gel was placed in an electrophoresis chamber and covered with TBE. 15 $\mu$L of samples containing 750 ng of DNA were loaded onto the gel along with a DNA reference ladder (GeneRuler Ultra Low Range, Thermo Scientific). A potential of 75 V (3.75 V cm$^{-1}$) was applied for 120 minutes. The gel was then imaged using a GelDoc-It system containing a UV lamp for illumination.

**Fluorescence Microscopy of GUVs:** Glass slides, cleaned with 15 min sonication cycles (Hellmanax III 2%/isopropanol/MilliQ water) were covered with a solution containing bovine serum albumin (BSA) at 0.1% (w/v) and placed in a pre-heated oven ($T > 60$°C) for an hour to allow for passivation. After thoroughly rinsing with milliQ water to remove excess BSA and dried, silicone incubation chambers (Sigma-Aldrich) were stuck to them and sealed with DNAse-free tape to prevent evaporation. For imaging, GUVs were carefully pipetted into the chambers and allowed to sink for at least 10 min before acquisition. Micrographs were recorded at 1400 Hz and averaging over 10 frames using a Leica TCS SP5 confocal microscope equipped with an HC PL APO CORR CS 40× / 0.85 dry objective from Leica. TexasRed (excitation maximum - 596 nm; emission maximum - 615 nm) was tracked using a He-Ne laser (594 nm) for excitation, and Alexa488/fluorescein (excitation maximum - 495 nm; emission maximum - 520 nm) signal was excited using an Ar-ion laser (488 nm).

**Data Acquisition:** Throughout this work, confocal micrographs of DNA-functionalized GUVs were used to assess the partitioning tendency of the nanostructures to programmably control their distribution and transport within model membranes. The screening and acquisition process followed simple guiding criteria, where individual GUVs were imaged when:

1. The vesicle was big enough such that Brownian motion was negligible within the acquisition time (of under a minute).

2. The vesicle was small enough to fit in a 64.84 $\mu$m × 64.84 $\mu$m field of view (the largest our confocal microscope can acquire at the fastest scanning rate of 1400 Hz).

3. The vesicle was oriented with the plane separating the two quasi-hemispherical domains roughly perpendicular to the imaging plane, which in turn results in a visible sharp interface between the two phases at the equatorial section of the GUV (as exemplified in the many micrographs presented in the manuscript).

While criterion 3) is solely related to vesicle orientation, and thus has no impact on the physics of lateral partitioning of nanostructures, the size cut-off introduced by criteria 1) and 2) could bias the measured partitioning observable if it were to depend on GUV size. The latter has been ruled out in Fig. S4, where the partitioning tendency is presented as a function of vesicle radius. In here, the lack of correlation between the two variables indicates that the natural polydispersity of electroformed GUVs does not influence the partitioning behaviour of the nanostructures, which consequently confirms that the size-related criteria for acquisition do not bias the results.

The emergence of wave-like patterns in the intensity profiles, such as the one observed for the lipid channel in Fig. S1c, follows from an artefact related to confocal-microscopy scanning. We observed that in those regions where the membrane is aligned with the scanning direction, the fluorescence intensity appeared higher compared to that detected on membrane regions oriented perpendicular to scanning. The result is that, for a uniformly-fluorescent GUV, an equatorial cross section would show two brighter regions on opposite sides, separated by two dimmer regions. In other words, fluorescence intensity completes two full oscillations around the circumference of a GUV's cross section. This artefact has little or no impact on our measurements aimed at detecting the average fluorescence intensity in quasi-hemispherical domains, owing to the fact that the detected intensity modulation completes one full oscillation over the half-circle associated to each domain in the equatorial cross section, and hence its effects average out. Additionally, the artefact appears mainly (or uniquely) on the TexasRed lipid channel, which used only for segmentation purposes, rather than on the DNA channel used to evaluate partitioning. Nonetheless, to definitely eliminate any possibility of bias, for instance related to the fact that the two domains may not be perfect half-spheres, each vesicle in this study was imaged twice, with a 90° difference in the scanning direction (which can be freely rotated with our instrument). The data from the two orientations were then averaged.

**Data Processing and Analysis:** The image analysis, as detailed in Fig. S1, is based on the user-assisted segmentation of the membrane radial intensities and their subsequent classification to either the $L_{\mathrm{o}}$ or $L_{\mathrm{d}}$ phases. When performing image analysis, micrographs were only discarded when bright lipid aggregates were present in the membrane due to imperfect electroformation, as they would make it challenging for the software to distinguish the $L_{\mathrm{o}}$ from the $L_{\mathrm{d}}$ domains, and bias the estimates of DNA nanostructure fluorescence.

Briefly, the custom-built MATLAB software loads each set of micrographs for the DNA and Lipid channels, and processes the images for with a Gaussian (blurring) filter to remove noise and facilitate segmentation. Through a Graphical User Interface, rough positions for the center and radius of the vesicle are manually selected. These estimated values are then used to enclose the membrane contour within a ring, as shown graphically in Fig. S1a. The software subsequently extracts 361 equally-spaced radial profiles within the defined ring and, for each, locates the position of the membrane through a Gaussian fit (see Fig. S1b). The extracted membrane positions are then used to acquire, from the un-filtered (raw) images, the average fluorescence intensity of the membrane through numerical integration around a width $w = \pm 2$ px from the fitted centre of the Gaussian. The average fluorescence intensity values are later used to construct an intensity profile around the circumference of the equatorial section of membrane, as exemplified in Fig. S1c, for both the signal acquired in the DNA and Lipid channels. Afterwards, the Lipid profile (red in Fig. S1c) was used as a reference to threshold and identify the angular coordinates of the dark domains ($L_{\mathrm{d}}$), and thus classify the DNA intensities to either of the $L_{\mathrm{o}}$ or $L_{\mathrm{d}}$ phases.

In some instances, where the partitioning of DNA nanostructures resulted in dark domains with intensities close or similar to that of the background, the segmentation algorithm described above and presented in Fig. S1 was unable to accurately locate the radial position of the membrane. This issue was particularly severe for sT constructs at the lowest DNA/lipid molar ratios in Fig. S7. This type of artefact, exemplified with a representative micrograph in Fig. S2a, required additional processing and an improved segmentation algorithm to extract reliable partitioning metrics. The correct identification of membrane location was achieved by applying a circle fitting routine. We considered a sub-set of the radial-profile points on the membranes initially identified by the algorithm, namely those within $\pm$ 5% of the estimated average radius of the vesicles. These points were fitted to a circle, which was then used as a better estimate of the membrane location and onto which fluorescence intensity was evaluated, as shown in Fig. S2b. This algorithm allowed to correct for imperfect segmentation as required.

Finally, the $L_\mathrm{o}$ and $L_\mathrm{d}$-associated DNA intensities ($I_{L_\mathrm{o}}$ and $I_{L_\mathrm{d}}$, respectively), extracted from micrographs as detailed above, were used to calculate the fractional intensity $f_{\mathrm{p},L_\mathrm{o}} = I_{L_\mathrm{o}}/\left(I_{L_\mathrm{o}} + I_{L_\mathrm{d}}\right)$, which we used as observable to describe the partitioning behavior of DNA nanostructures in the membrane. In that sense, the data from each pair of micrographs associated to each vesicle (imaged with the 90° difference in scanning direction) were processed independently and ultimately averaged to provide a single $f_{\mathrm{p},L_\mathrm{o}}$ value describing each vesicle.

### Supplementary Discussion 1: Impact of fluorescence cross-talk and FRET.

The fluorescent lipid TexasRed-DHPE was used as a marker to identify $L_\mathrm{d}$ domains due to its well characterized partitioning preference and photo-stability. However its presence may in principle bias the partitioning observable $f_{\mathrm{p},L_\mathrm{o}}$ through two avenues: cross-talk between fluorescence channels and Förster Resonance Energy Transfer (FRET).

While cross-talk most prominently occurs from shorter to longer wavelength channels (i.e., Alexa488 (green, DNA) to TexasRed (red, lipid) channels), the TexasRed signal is not used for quantitative assessment of the partitioning tendency, but only used as a reference to identify the lipid phases (see Fig. S1). Owing to the strong partitioning of TexasRed-DHPE in $L_\mathrm{d}$, green-to-red cross-talk has no impact on the ability of the analysis algorithm to accurately identify lipid phases.

In turn, the much less substantial red-to-green cross-talk, from the lipid/TexasRed channel to the DNA/Alexa488 channel, may bias the determination of the Alexa488 intensities, and thus the resulting $f_{\mathrm{p},L_\mathrm{o}}$.

The degree of bleed from red to green channels must be proportional to the concentration of the TexasRed dye, and therefore, to the fluorescence intensity recorded in the red (lipid) channel. Therefore, a correction could be applied for red-to-green cross-talk using

$$I_{L_\mathrm{x},\mathrm{corr}} = I_{L_\mathrm{x}} - A \times I_{L_\mathrm{x},\mathrm{TXR}}, \tag{S1}$$

where $I_{L_\mathrm{x}}$ is the fluroescence intensity recorded in the green (DNA) channel, $I_{L_\mathrm{x},\mathrm{TXR}}$ the one recorded in the red channel, and $I_{L_\mathrm{x},\mathrm{corr}}$ the corrected DNA signal. The label $L_\mathrm{x}$ indicates that the fluorescence is measured either in the $L_\mathrm{o}$ or $L_\mathrm{d}$ lipid phases.

The correction factor $A$ in Eq. S1 was determined as the ratio between $I_{L_\mathrm{x}}$ and $I_{L_\mathrm{x},\mathrm{TXR}}$ as recorded in a population of GUVs containing only the TexasRed-labelled lipids, but not the DNA nanostructures, in the same settings as the ones used for data acquisition. These measurements provided $A = 0.03 \pm 0.005$.

When Eq. S1 is applied to $f_{\mathrm{p},L_\mathrm{o}} = I_{L_\mathrm{o}}/\left(I_{L_\mathrm{o}} + I_{L_\mathrm{d}}\right)$, the correction gives

$$f_{\mathrm{p},L_\mathrm{o}} = \frac{I_{L_\mathrm{o}} - A \times I_{L_\mathrm{o},\mathrm{TXR}}}{I_{L_\mathrm{o}} - A \times I_{L_\mathrm{o},\mathrm{TXR}} + I_{L_\mathrm{d}} - A \times I_{L_\mathrm{d},\mathrm{TXR}}} = \frac{I_{L_\mathrm{o}}}{I_{L_\mathrm{o}} + I_{L_\mathrm{d}}\left(\frac{1-A\times I_{L_\mathrm{d},\mathrm{TXR}}/I_{L_\mathrm{d}}}{1-A\times I_{L_\mathrm{o},\mathrm{TXR}}/I_{L_\mathrm{o}}}\right)}, \tag{S2}$$

Given the small magnitude of $A$, the correction in Eq. S2 is also small, as demonstrated in Fig. S3a with the comparison between the corrected and non-corrected $f_{\mathrm{p},L_\mathrm{o}}$ values for sT and dC constructs.

In all cases, and particularly for sT, the shift on the median of the distributions caused by the cross-talk correction is small compared to the intrinsic statistical variability of $f_{\mathrm{p},L_\mathrm{o}}$, and thus carries no impact on the conclusions of the work presented here. In addition, it can be observed that, particularly for the case of dC, the correction produces a substantial broadening of the distribution, likely caused by the additional stochastic uncertainties associated to $I_{L_\mathrm{d},\mathrm{TXR}}$ and $I_{L_\mathrm{o},\mathrm{TXR}}$, which appear in the corrected Eq. S2.

Owing to the negligible impact of the cross-talk correction and its detrimental effect on the width of the distribution of the computed $f_{\mathrm{p},L_\mathrm{o}}$, we chose not to apply the correction to the data presented throughout this work. This choice is further justified by the direct comparison of the $f_{\mathrm{p},L_\mathrm{o}}$ values with those measured in the absence of the TexasRed lipid, and hence cross-talk, as collated in Fig. S3a, which show good alignment with the uncorrected data.

Let us now examine the potential impact of FRET between the DNA-bound Alexa488 (donor) and lipid-bound TexasRed (acceptor). If FRET were to take place with efficiency $\phi_{\mathrm{FRET},L_\mathrm{x}}$, the DNA fluorescence intensity could be corrected as

$$I_{L_\mathrm{x},\mathrm{corr}} = \frac{I_{L_\mathrm{x}}}{1 - \phi_{\mathrm{FRET},L_\mathrm{x}}}, \tag{S3}$$

which applied to $f_{\mathrm{p},L_\mathrm{o}}$ gives

$$f_{\mathrm{p},L_\mathrm{o},\mathrm{corr}} = \frac{I_{L_\mathrm{o}}}{I_{L_\mathrm{o}} + I_{L_\mathrm{d}}\left(\frac{1-\phi_{\mathrm{FRET},L_\mathrm{o}}}{1-\phi_{\mathrm{FRET},L_\mathrm{d}}}\right)}, \tag{S4}$$

Importantly, Eq. S4 shows that FRET would only affect the measurements if the efficiency were to be different in the different lipid phases, *i.e.* $\phi_{\mathrm{FRET},L_\mathrm{d}} \neq \phi_{\mathrm{FRET},L_\mathrm{o}}$.

The FRET efficiencies, summarized in Fig. S3b, were experimentally determined using $\phi_{\mathrm{FRET},L_\mathrm{x}} = 1 - \frac{I_{L_\mathrm{x}}}{I^{\mathrm{NoTXR}}_{L_\mathrm{x}}}$ where, as usual, $I_{L_\mathrm{x}}$ is the intensity of DNA-labelled Alexa488 (donor) recorded on GUVs that feature the fluorescent TXR-lipids (acceptor), and $I^{\mathrm{NoTXR}}_{L_\mathrm{x}}$ is the same quantity, but recorded for GUVs that lack TexasRed lipids. It can be readily observed in Fig. S3 that the estimates of $\phi_{\mathrm{FRET},L_\mathrm{x}}$ carry very large error bars, which are ascribable to the large statistical fluctuations in the fluorescence intensities among populations of GUVs and the propagation of the resulting uncertainties. Although well aware of the limited interpretability of these results, it is interesting to highlight that, while FRET efficiency is small (or zero) for dC-bearing duplexes, it appears to be non-zero for sT anchors. This difference may follow from the fact that cholesterol and tocopherol moieties could insert within the bilayer at different depths, hence resulting in a different distance between the lipid and DNA-bound dyes. Nonetheless, for both sT and dC, $\phi_{\mathrm{FRET},L_\mathrm{d}} \sim \phi_{\mathrm{FRET},L_\mathrm{o}}$ suggesting that the correction in Eq. S4 has no impact on the computed $f_{\mathrm{p},L_\mathrm{o}}$.

Similarly to the case of the cross-talk correction, we thus decided not to correct our data for the potential effect of FRET, owing to the negligible impact and the very large uncertainties associated with propagating the exceedingly large $\phi_{\mathrm{FRET},L_\mathrm{x}}$ error to $f_{\mathrm{p},L_\mathrm{o}}$.

Ultimately, the negligible impact of cross-talk and FRET is demonstrated by the control experiment carried out on vesicles without the fluorescent lipid (Fig. S3a), which demonstrates that $f_{\mathrm{p},L_\mathrm{o}}$ does not depend on the presence or absence of the TexasRed labelled lipids.

Note that for some DNA constructs, namely nanostars dC/sT (in Fig. 3) and the duplexes in Fig. S6, a fluorescein tag was used instead of Alexa488. The similar spectral characteristics of the two dyes imply that the controls and arguments discussed in this section apply to fluorescein-labelled constructs as well.

### Supplementary Figures

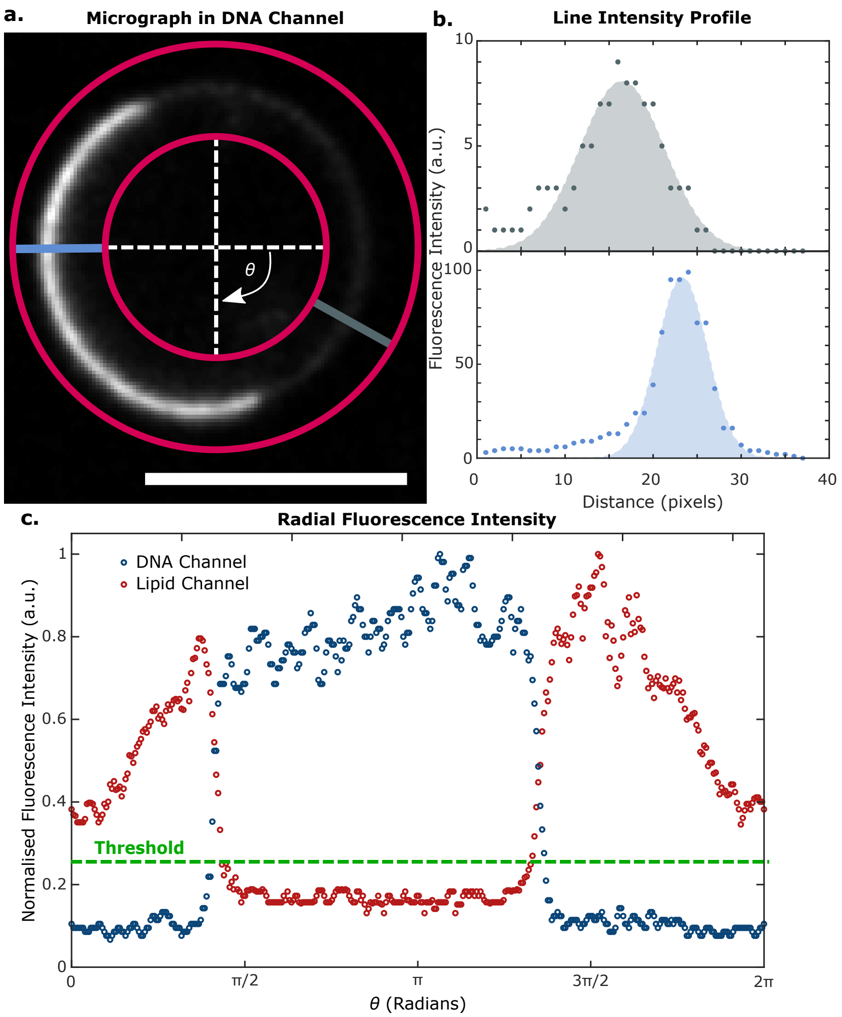


**Figure S1: Image analysis pipeline for measuring DNA-construct fluorescence intensity in equatorial confocal micrographs.** Equatorial confocal micrographs, acquired simultaneously for the lipid (TexasRed) and DNA (Alexa488 or fluorescein) channels, are used as input for our custom-built analysis pipeline. **a.** A line connecting two concentric circles, which enclose the membrane circumference within a ring, is projected radially using polar coordinates from $\theta = 0$ to $2\pi$ to acquire intensity profiles in steps of $\pi/180$. **b.** From individual radial profiles we extract the average intensity *via* numerical integration around a width ($w = \pm 2$ px) from the membrane peak, which is in turn determined with a Gaussian fit. **c.** These intensity values are used to construct an intensity profile around the circumference of the vesicle equatorial section for both fluorescent channels. The $L_\mathrm{o}$ and $L_\mathrm{d}$ domains are determined *via* thresholding of the circumference profile acquired in the lipid channel (TexasRed). Subsequently, the DNA fluorescent signal is averaged over the angles identified as belonging to $L_\mathrm{o}$ and $L_\mathrm{d}$ to extract $I_{L_\mathrm{o}}$ and $I_{L_\mathrm{d}}$, which are then is used to calculate a fractional intensity in $L_\mathrm{o}$ ($f_{p,L_\mathrm{o}}$). Scale bar = 10 $\mu$m.

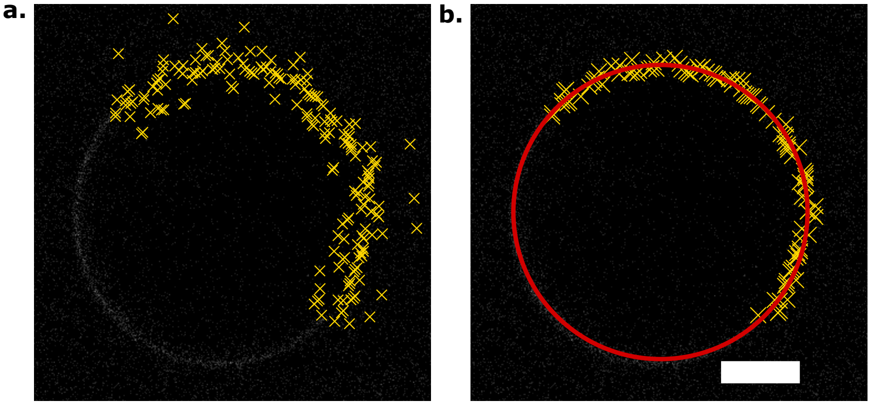


**Figure S2: Circle fitting routine to correct for imperfect segmentation of low-intensity equatorial membrane domains.** DNA nanostructure partitioning can result in dark (low intensities) domains, in the order of the background noise. The latter leads to segmentation imperfections when processed with the analysis module described in Fig. S1. **a.** Micrograph with the initially identified membrane positions (yellow), showcasing the deviation due to noise. **b.** A second segmentation module processes a subset of the points identified in **a**, namely those that are within ±5% of the estimated radius of the vesicle. These points, shown in panel **b**, are subjected to a circle fitting routine, and the average membrane intensity values are extracted at a distance equal to the radius of the circle (in red). Scale bar = 10 $\mu$m.

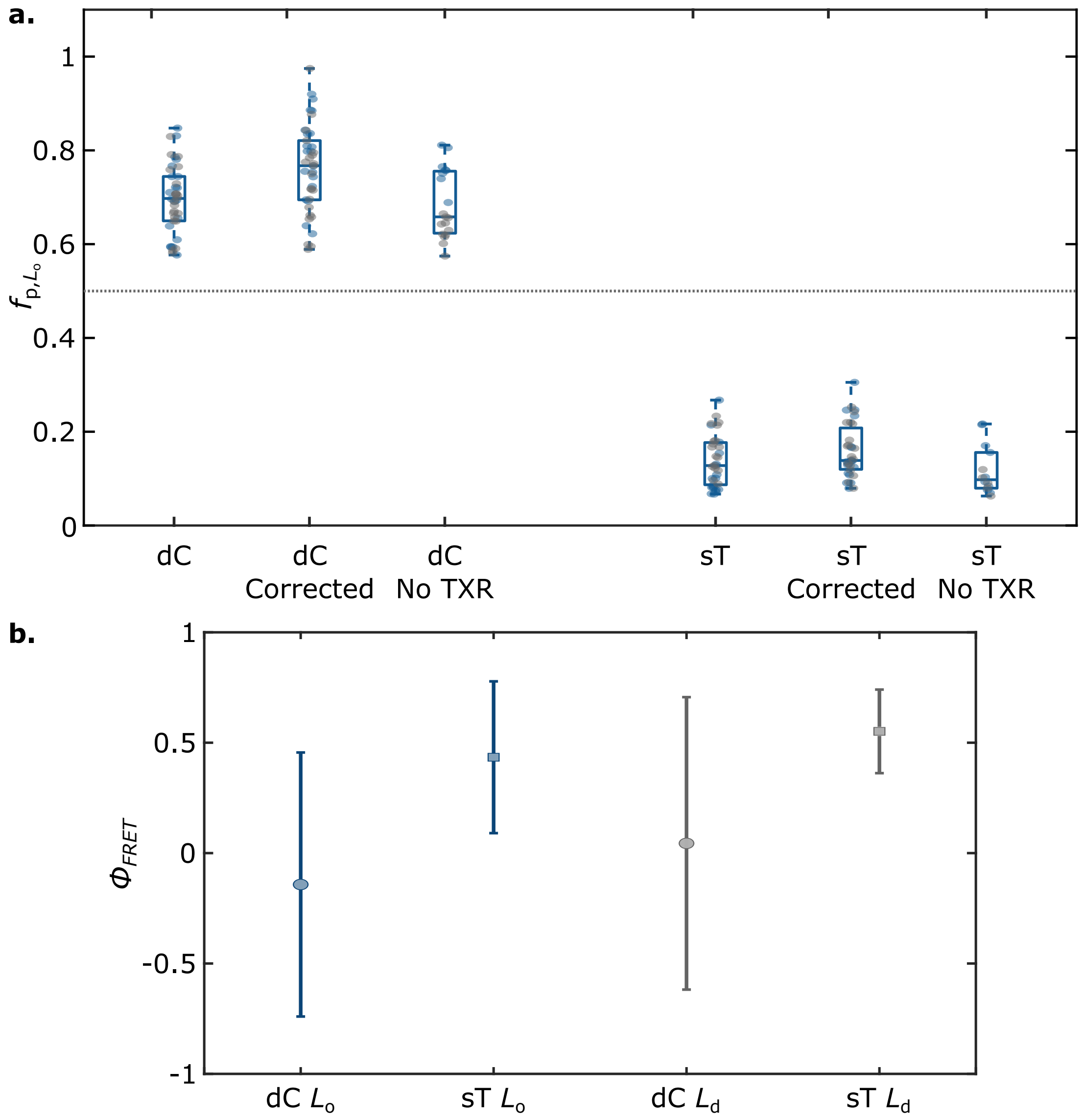


**Figure S3: The presence of the fluorescent lipid marker has a negligible impact on the quantitative description of partitioning tendency of nanostructures.** **a.** Fractional intensity in $L_o$ ($f_{p,L_o}$) of two DNA nanostructures (dC and sT), conveyed by box-scatter plots, for two individual repeats (blue/grey, respectively) recorded on ternary GUV populations in the presence and absence of TexasRed. In the presence of the fluorescent lipid, we also show the data as processed using Eq. S2 to correct for fluorescence cross-talk (see Supplementary Discussion 1). For the case of GUVs lacking fluorescent lipids, the thresholding step to identify $L_o$ and $L_d$ domains was carried out using the fluorescent traces of the DNA channel. Results show no difference in $f_{p,L_o}$ for GUVs with and without TexasRed. The dotted line indicates no partitioning ($f_{p,L_o} = 0.5$). **b.** Computed FRET efficiencies $\phi_{FRET}$ for dC (circles) and sT (squares) in both $L_o$ (blue) and $L_d$ (grey) phases. In all cases, $\phi_{FRET,L_d} \sim \phi_{FRET,L_o}$, indicating a negligible effect on the extracted $f_{p,L_o}$ (Eq. S4). See Supplementary Discussion 1 for information of FRET efficiency measurements.

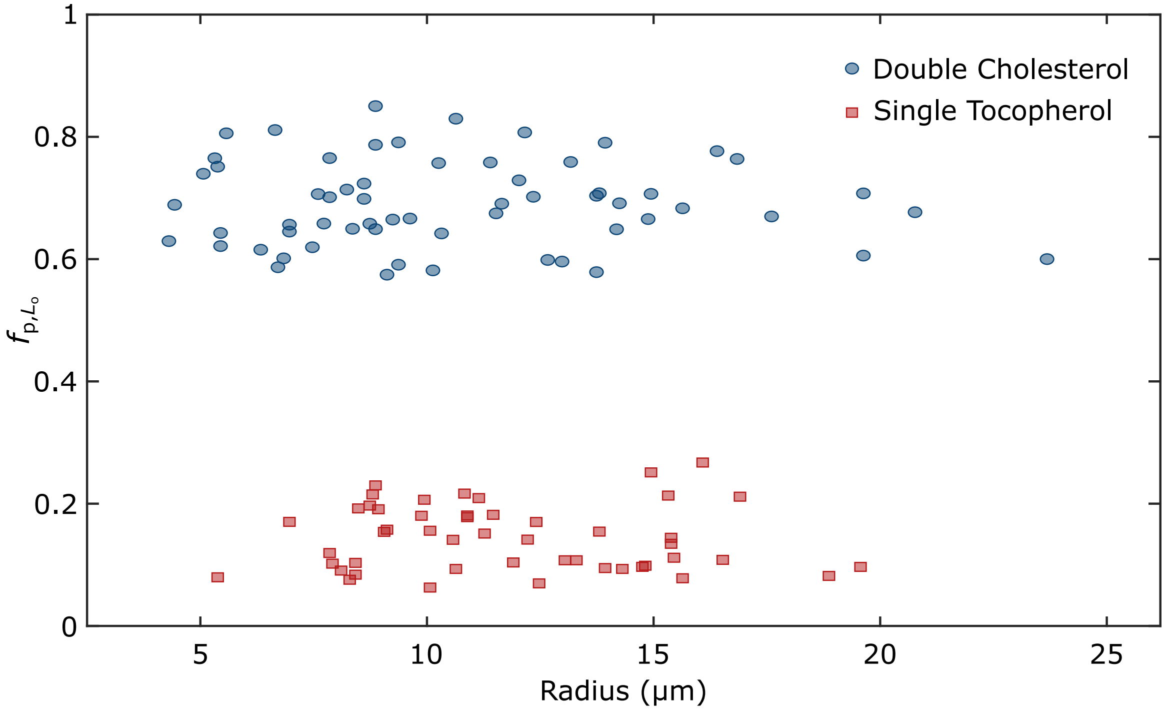


**Figure S4: GUV polydispersity does not influence DNA nanostructure partitioning.** Fractional intensity in $L_o$ ($f_{p,L_o}$) of DNA nanostructures anchored *via* dC (blue circles) or sT (red squares) as a function of vesicle radius, recorded on ternary GUV populations. The lack of correlation between the two variables indicates that the natural size heterogeneity of electroformed vesicles does not have an effect on the lateral distribution of DNA nanostructures.

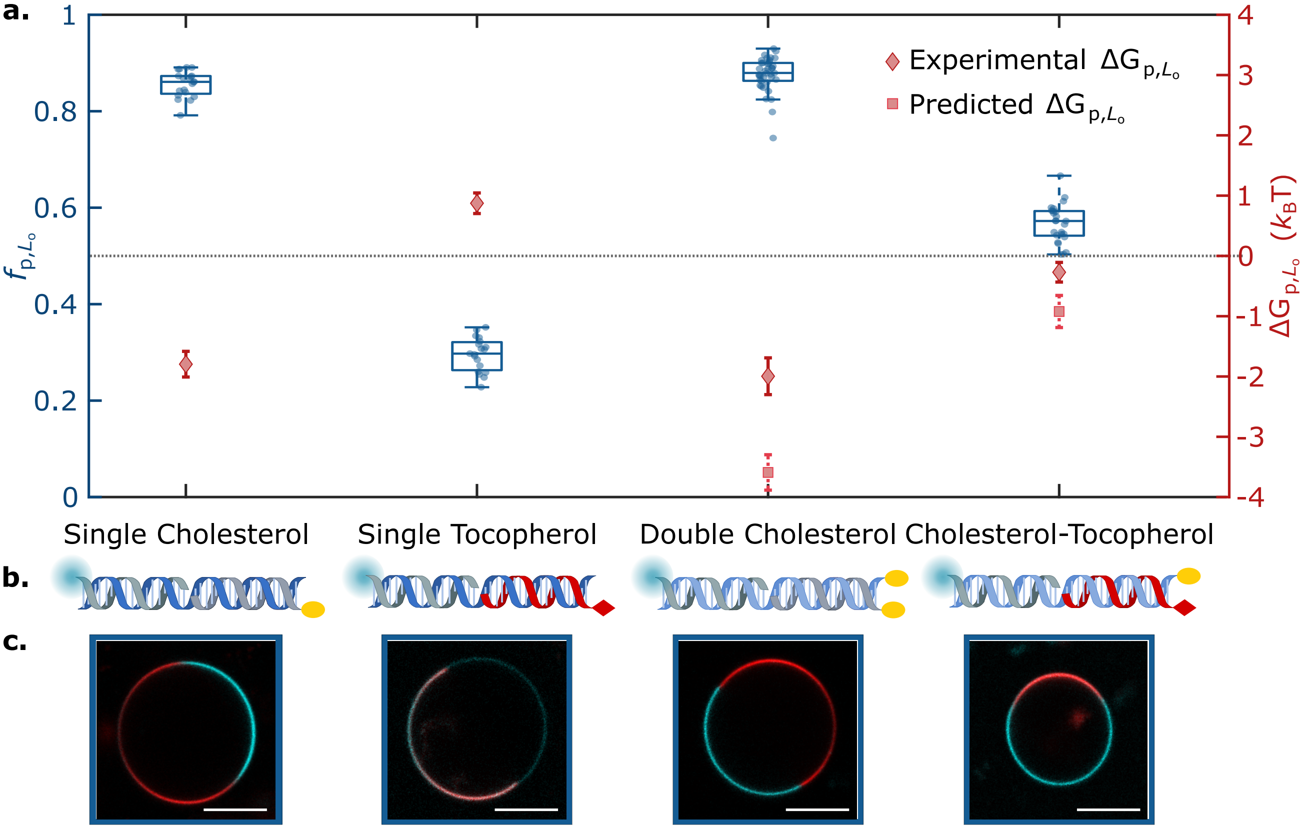


**Figure S5: Lateral distribution of duplex DNA nanostructures in quaternary (DOPC/DPPC/cardiolipin/chol) GUVs.** Data are analogous to those shown in Fig. 2 for the case of ternary GUVs. **a.** Fractional intensity ($f_{\mathrm{p},L_\mathrm{o}}$) and free energy of partitioning ($\Delta G_{\mathrm{p},L_\mathrm{o}}$) of DNA nanostructures in $L_\mathrm{o}$, conveyed by box-scatter plots and lozenges respectively, for DNA-decorated quaternary GUV populations featuring different anchoring motifs: single cholesterol-TEG (sC), double cholesterol-TEG (dC), single tocopherol (sT), or a combination of single cholesterol-TEG and single tocopherol (sC+sT). For dC and sC+sT, squares indicate the predicted free energy values for combined anchors, determined from Eq. 1 (main text) using measured values of $\Delta G^{\mathrm{sT}}_{\mathrm{p},L_\mathrm{o}}$ and $\Delta G^{\mathrm{sC}}_{\mathrm{p},L_\mathrm{o}}$. The dotted line indicates no partitioning ($f_{\mathrm{p},L_\mathrm{o}} = 0.5$, $\Delta G_{\mathrm{p},L_\mathrm{o}} = 0$). **b.** Schematic depictions of DNA duplexes bearing sC, sT, dC or sC+sT. **c.** Representative confocal micrographs of the DNA-functionalized GUVs. The $L_\mathrm{d}$ phase is labelled with TexasRed (red), and DNA constructs (cyan) with Alexa488. Scale bars = 10 $\mu$m.

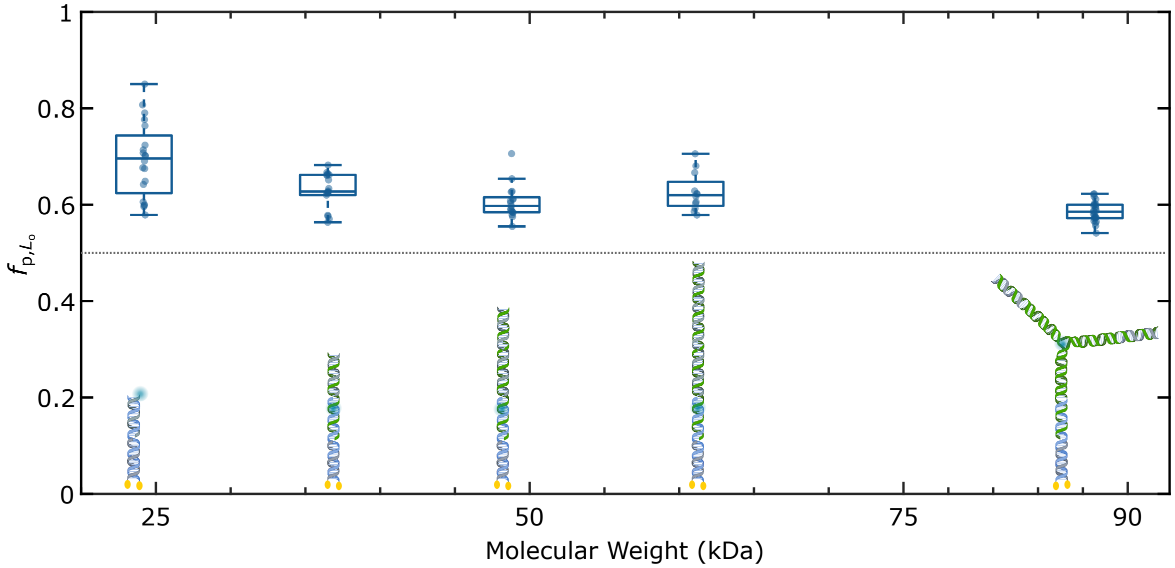


**Figure S6: Lateral distribution of DNA nanostructures of increasing molecular weight.** Fractional intensity ($f_{p,L_o}$) of DNA nanostructures in $L_o$, conveyed by box-scatter plots, for GUV populations decorated with duplexes of increasing molecular weight, and compared to the DNA nanostar (schematically depicted in scale). The dotted line indicates no partitioning ($f_{p,L_o} = 0.5$). The increase of size and molecular weight impacts the partitioning, possibly owing to steric encumbrance or electrostatic repulsion limiting the accumulation in $L_o$.

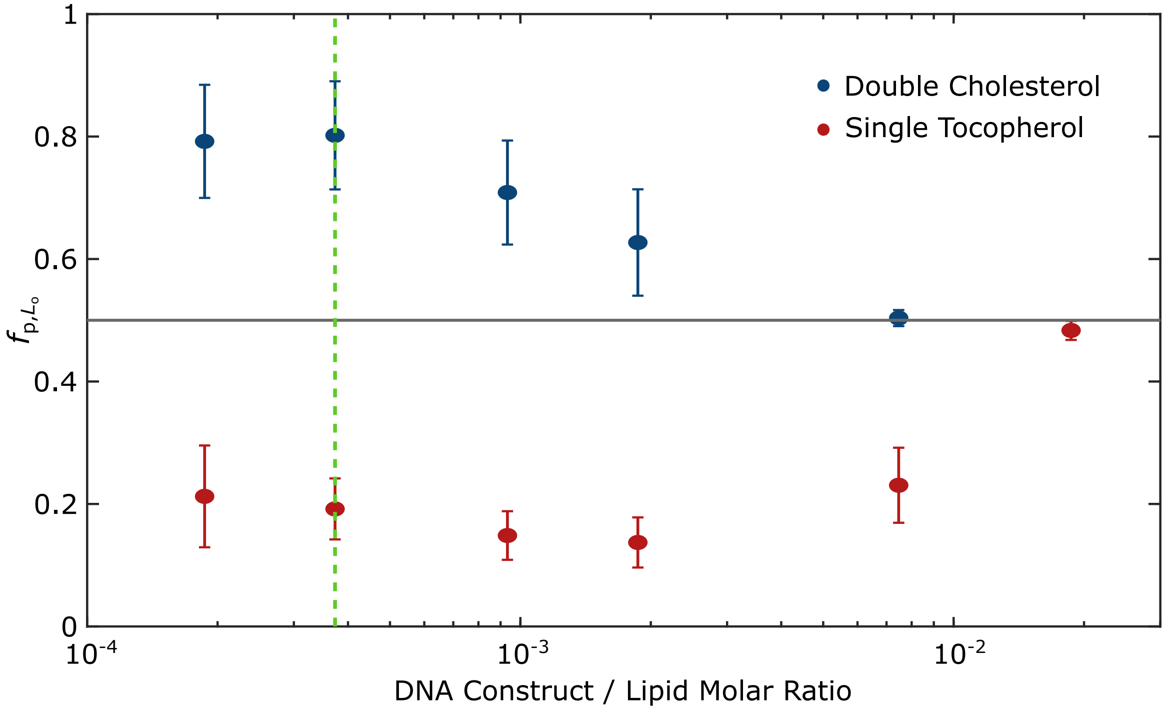


**Figure S7: High DNA/lipid ratio suppresses partitioning due to membrane saturation.** Fractional intensity ($f_{p,L_o}$) of DNA nanostructures in $L_o$, conveyed by plots of means ± standard deviations, for GUV populations decorated with DNA duplexes (dC in blue; sT in red) at increasing nominal DNA/Lipid molar ratios, and thus resulting in different densities of DNA constructs within the bilayers. The molarity of GUVs was determined using a phosphatidylcholine (PC) enzymatic assay. The dotted line indicates no partitioning ($f_{p,L_o} = 0.5$). The green dash line marks the DNA/lipid ratio ($\sim 4 \times 10^{-4}$) used throughout this work. The suppression of partitioning at high DNA/lipid ratios further supports the hypothesis that steric interactions between the nanostructures modulate the lateral distribution to a point where accumulation in a given phase leads to their saturation and subsequent impossibility to accommodate more constructs. Membrane saturation occurs at higher DNA/lipid ratios for the case of sT compared to dC. We ascribe this shift to a lower overall membrane affinity of the sT constructs, which may result in a lower density of membrane-anchored motifs at fixed DNA/lipid ratios.

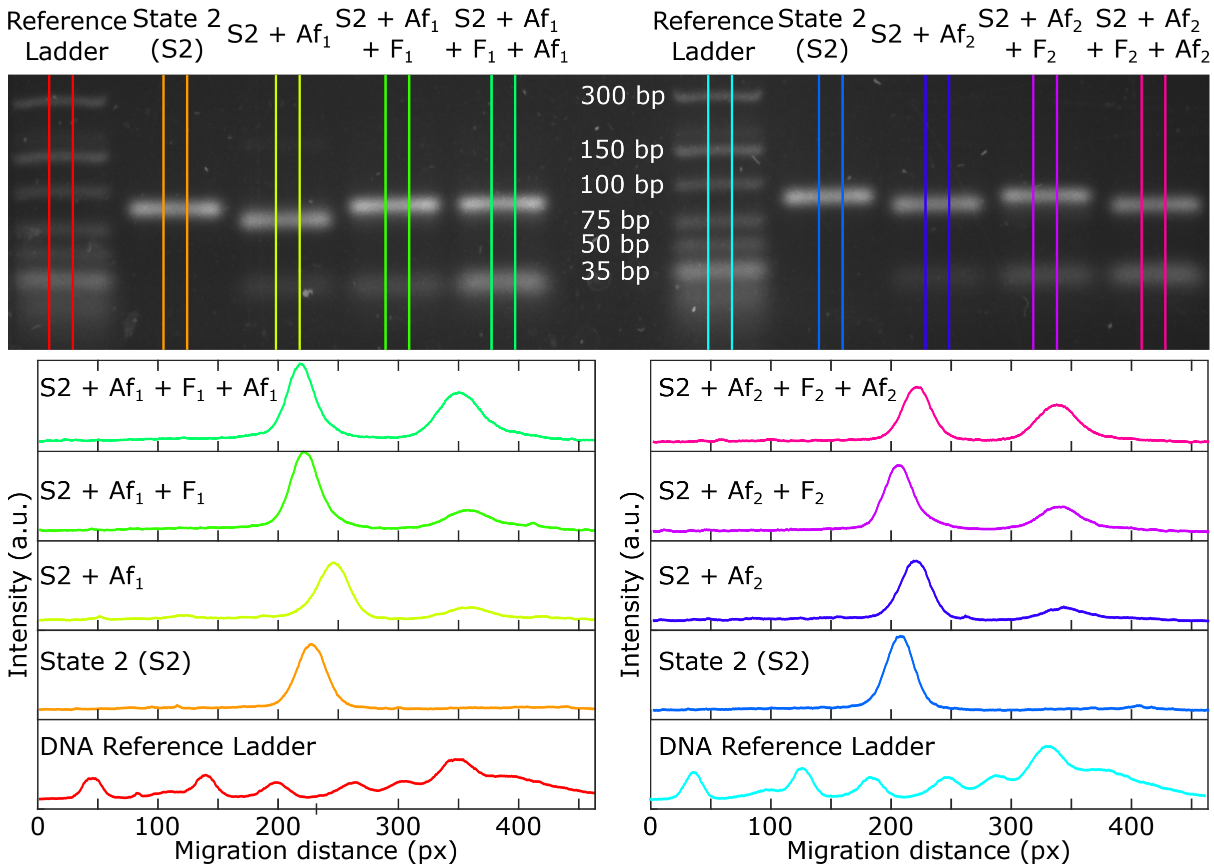


**Figure S8: DNA nanostructure reconfigurability *via* toeholding reactions demonstrated by AGE.** Agarose Gel Electrophoresis (AGE) of constructs described in Fig. 4 (without hydrophobic modifications) showing the functionality of toehold domains ($\alpha_2$ and $\delta_2$, left; $\alpha_1$ and $\delta_1$, right). State 2 (S2) constructs were incubated with combinations of $\text{Antifuel}_{1(2)}$ ($\text{Af}_{1(2)}$) and $\text{Fuel}_{1(2)}$ ($\text{F}_{1(2)}$) strands, as indicated in each lane and intensity profile, to probe the reconfiguration process. Each Antifuel/Fuel strand was added sequentially, with waiting times of $\sim$15 min before the subsequent addition, in 0.5$\times$ excess with respect to its prior step (e.g. S2 (1$\times$) + $\text{Af}_1$(1.5$\times$) + $\text{F}_1$(2$\times$)) + $\text{Af}_1$(2.5$\times$)). Expectedly, bands show different migration distances in the presence of Antifuel/Fuel, confirming nanostructure reconfiguration. The emergence of a bright band, just below the reference marker corresponding to 35 bp, points to the accumulation of waste (Fuel/Antifuel pairs). Importantly, reversibility is attainable at least for one cycle. While the $\alpha_1/\delta_1$ (right) pair of toeholds achieve two cycles of reconfiguration, $\alpha_2/\delta_2$ do not show the last reaction going to completion (at least within the experimental timescale $\sim$3 hrs). The latter suggests the presence of partial inefficiencies and/or small thermodynamic unbalances in the system.

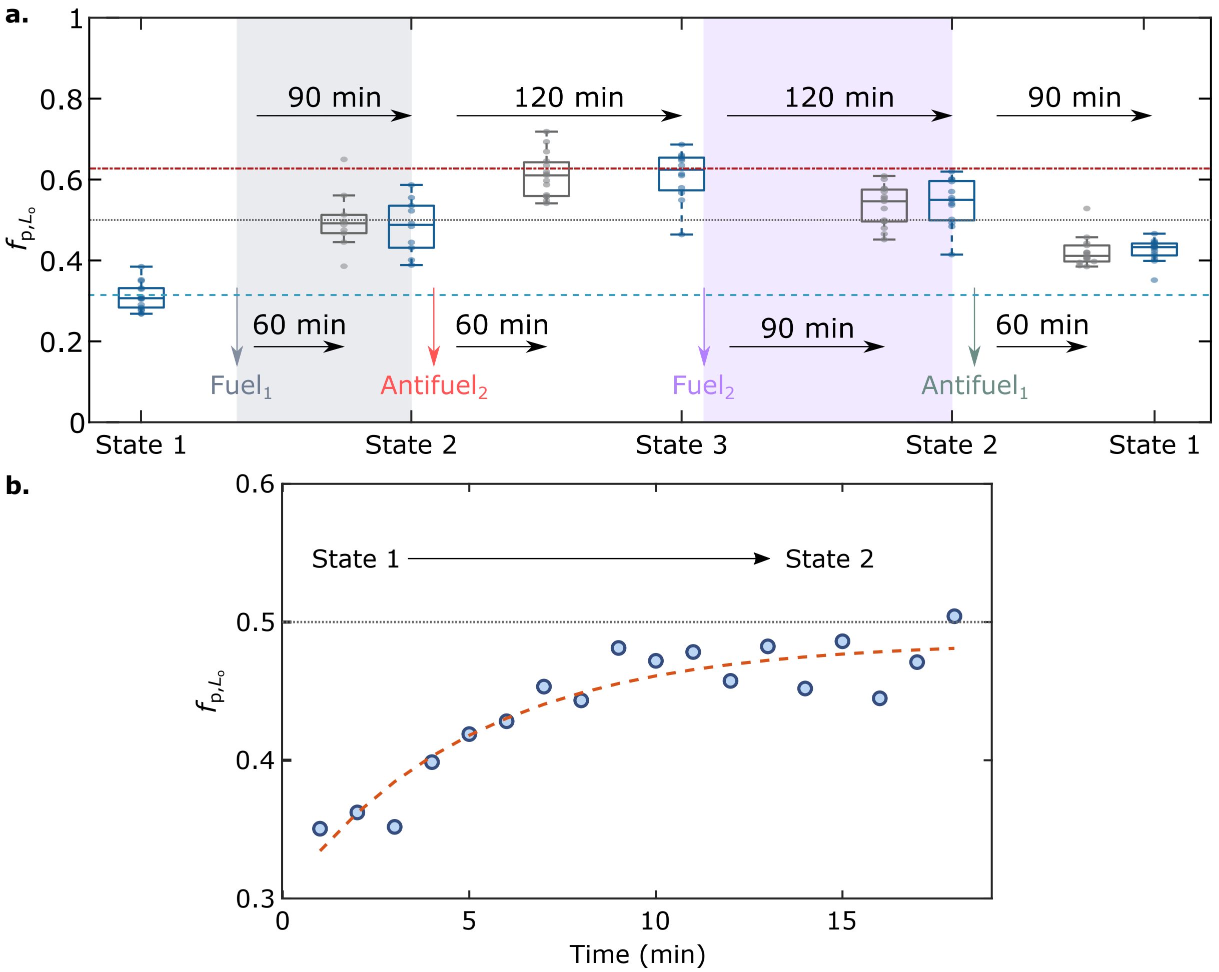


**Figure S9: DNA nanostructure reconfiguration and re-distribution reaches equilibrium in hundreds of seconds. a.** Data for the lateral re-distribution of the fluorescent cargo described in Fig. 4 at intermediate time points after the addition of fuel/antifuel (grey), which verify that the system reached equilibrium before the time points shown in Fig. 4b (blue). **b.** Time-evolution of $f_{\mathrm{p},L_\mathrm{o}}$ for an individual GUV undergoing lateral re-organisation of the fluorescent cargo in our responsive nanodevice upon adding Fuel$_1$, which switches the nanostructures from State 1 to State 2. The orange dashed line is an empirical fit to an exponential of the form $A - B \cdot \exp(-t/\tau)$, and the dotted line indicates no partitioning ($f_{\mathrm{p},L_\mathrm{o}}$ $= 0.5$). The fitted characteristic re-distribution time is $\tau \sim 5 \pm 3$ minutes. Three possible processes may limit construct re-distribution kinetics: i) the diffusion time of fuel/antifuel strands in the bulk, as they are gently added at the surface of the imaging chamber, ii) the rate of the strand displacement reactions, and iii) the diffusion time for re-distribution of membrane-anchored constructs. Considering the relevant length scales (mm) and the diffusion coefficient of free oligonucleotides at room temperature,[6] we expect the time-scales of i) to be in the order of hundreds to thousands of seconds. Process ii), given the concentrations of fuel/antifuel (100-300 nM) and the typical second-order rate constants for toeholding ($\sim$ $10^6$ $\mathrm{M}^{-1}$ $\mathrm{s}^{-1}$[7,8]), should occur over tens to hundreds of seconds, while iii) should take tens of seconds in light of the diffusion coefficient of membrane-anchored DNA nanostructures.[9,10] We can thus conclude that i) is most likely the rate-limiting process in our experiments, compatibly with the measured relaxation timescales of $\sim 5$ minutes

### DNA constructs and sequences

**Table S1: Nanostructures and oligonucleotides that comprise them.** Each construct is self-assembled through a quenching temperature ramp as discussed in the Methods at a final concentration of 2 $\mu$M and then diluted for GUV functionalization. Constructs require strands in 1:1 stochiometric ratios, unless stated otherwise in the case of the nanostars.

| **Construct** | **Strands** |
|---|---|
| $\mathrm{Duplex}_{\mathrm{sC}}$ | $\mathrm{B}_{\mathrm{bb,chol}} + \mathrm{B}_{\mathrm{b}} + \mathrm{Mem}^{\mathrm{f}}$ |
| $\mathrm{Duplex}_{\mathrm{sT}}$ | $\mathrm{B}_{\mathrm{bb,toc}} + \mathrm{B}_{\mathrm{b}} + \mathrm{Mem}^{\mathrm{f}}$ |
| $\mathrm{Duplex}_{\mathrm{dC}}$ | $\mathrm{B}_{\mathrm{bb,chol}} + \mathrm{B}_{\mathrm{b,chol}} + \mathrm{Mem}^{\mathrm{f}}$ |
| $\mathrm{Duplex}_{\mathrm{sT+sC}}$ | $\mathrm{B}_{\mathrm{bb,toc}} + \mathrm{B}_{\mathrm{b,chol}} + \mathrm{Mem}^{\mathrm{f}}$ |
| $2\times$-$\mathrm{duplex}_{\mathrm{dC+dC}}$ | $\mathrm{B}_{\mathrm{bb,chol}} + \mathrm{B}_{\mathrm{b,chol}} + \mathrm{Lin}^{\mathrm{f},*} + \mathrm{C}_{\mathrm{b,chol}} + \mathrm{C}_{\mathrm{bb,chol}}$ |
| $2\times$-$\mathrm{duplex}_{\mathrm{sT+dC}}$ | $\mathrm{B}_{\mathrm{bb,toc}} + \mathrm{B}_{\mathrm{b}} + \mathrm{Lin}^{\mathrm{f},*} + \mathrm{LinF}_{6}\ \mathrm{C}_{\mathrm{b,chol}} + \mathrm{C}_{\mathrm{bb,chol}}$ |
| $\mathrm{Nanostar}_{\mathrm{dC}}$ | $\mathrm{D}_{\mathrm{bb,chol}} + \mathrm{D}_{\mathrm{b,chol}} + \mathrm{Arm}_1^{\mathrm{f}} + \mathrm{Arm}_2 + \mathrm{Arm}_{3,\mathrm{chol}} + (2\times)\mathrm{Arm}_{\mathrm{block,B_b}}$ |
| $\mathrm{Nanostar}_{\mathrm{sT}}$ | $\mathrm{B}_{\mathrm{bb,toc}} + \mathrm{B}_{\mathrm{b}} + \mathrm{Arm}_1^{\mathrm{f}} + \mathrm{Arm}_2 + \mathrm{Arm}_{3,\mathrm{toc}} + (2\times)\mathrm{Arm}_{\mathrm{block,D_b}}$ |
| State $1_{\mathrm{sT,cargo}}$ | $\mathrm{B}_{\mathrm{bb,toc}} + \mathrm{B}_{\mathrm{b}} + \mathrm{Lin}^{\mathrm{f},*} + \mathrm{LinF}_{12} + \mathrm{Antifuel}_1$ |
| State $1_{\mathrm{dC}}$ | $\mathrm{C}_{\mathrm{b,chol}} + \mathrm{C}_{\mathrm{bb,chol}}$ |
| State $2_{\mathrm{sT,cargo,dC}}$ | $\mathrm{B}_{\mathrm{bb,toc}} + \mathrm{B}_{\mathrm{b}} + \mathrm{Lin}^{\mathrm{f},*} + \mathrm{LinF}_{12} + \mathrm{C}_{\mathrm{b,chol}} + \mathrm{C}_{\mathrm{bb,chol}}$ |
| State $3_{\mathrm{cargo,dC}}$ | $\mathrm{Lin}^{\mathrm{f},*} + \mathrm{LinF}_{12} + \mathrm{C}_{\mathrm{b,chol}} + \mathrm{C}_{\mathrm{bb,chol}} + \mathrm{Antifuel}_2$ |
| $\mathrm{Extended}_{1\mathrm{X}}$ $\mathrm{Duplex}_{\mathrm{dC}}$ | $\mathrm{D}_{\mathrm{bb,chol}} + \mathrm{D}_{\mathrm{b,chol}}$ + A1 + A1* |
| $\mathrm{Extended}_{2\mathrm{X}}$ $\mathrm{Duplex}_{\mathrm{dC}}$ | $\mathrm{D}_{\mathrm{bb,chol}} + \mathrm{D}_{\mathrm{b,chol}}$ + A1 + A1*-A2 + A2* |
| $\mathrm{Extended}_{3\mathrm{X}}$ $\mathrm{Duplex}_{\mathrm{dC}}$ | $\mathrm{D}_{\mathrm{bb,chol}} + \mathrm{D}_{\mathrm{b,chol}}$ + A1 + A1*-A2 + A2*-A3 + A3* |

**Table S2: Sequences of oligonucleotide strands.** Abbreviations: TEG: triethyleneglycol; FluorT: internal fluorescein modification on thymine (T).

| **Strand** | **Sequence (5' → 3')** |
|---|---|
| $\mathrm{B}_{\mathrm{bb,chol}}$ | /Cholesteryl-TEG/GTTGTGGTGGTGAGTGTG |
| $\mathrm{B}_{\mathrm{b}}$ | CATCTCACTACTCAACACCACACTCACCACCACAAC |
| $\mathrm{Mem}^{\mathrm{f}}$ | GTGTTGAGTAGTGAGATG/AlexaFluor488/ |
| $\mathrm{B}_{\mathrm{bb,toc}}$ | /Octyl-Tocopherol/GTTGTGGTGGTGAGTGTG |
| $\mathrm{B}_{\mathrm{b,chol}}$ | CATCTCACTACTCAACACCACACTCACCACCACAAC/Cholesterol-TEG/ |
| $\mathrm{C}_{\mathrm{b,chol}}$ | /Cholesteryl-TEG/CAATCACACCACAAACACCCAACACAACAACAAACC |
| $\mathrm{C}_{\mathrm{bb,chol}}$ | GTGTTTGTGGTGTGATTG/Cholesterol-TEG/ |
| $\mathrm{Lin}^{\mathrm{f},*}$ | GCTCTCTCATCACTAC/AlexaFluor488/ |
| $\mathrm{Lin}_6$ | GTGTTGAGTAGTGAGATGTTTGTAGTGATGAGAGAGCTTTGGTTTGTTGTTGTGTTGG |
| $\mathrm{LinF}_{12}$ | GTGTTGAGTAGTGAGATGATTCGCGTAGTGATGAGAGAGCGTTGTAGGTTTGTTGTTGTG |
| $\mathrm{Fuel}_1$ | GTTGTTGTGTTGGCCTTACTTCACG |
| $\mathrm{Fuel}_2$ | GTTTGAGGTGAAGTGTTGAGTAGTG |
| $\mathrm{Antifuel}_1$ | CGTGAAGTAAGGCCAACACAACAACAAACCTACAAC |
| $\mathrm{Antifuel}_2$ | GCGAATCATCTCACTACTCAACACTTCACCTCAAAC |
| $\mathrm{D}_{\mathrm{b,chol}}$ | CCAACACAACAACAAACCCGTTCCGACATAGAACCG/Cholesterol-TEG/ |
| $\mathrm{D}_{\mathrm{bb,chol}}$ | /Cholesteryl-TEG/ CGGTTCTATGTCGGAACG |
| $\mathrm{Arm}_1^{\mathrm{f}}$ | GTGTTGAGTAGTGAGATGTTGCAACGACTCCGCACTCGCGT/iFluorT/TGCGAATAGGTCTGTCAGTTT |
| $\mathrm{Arm}_2$ | GGTTTGTTGTTGTGTTGGTTAAACTGACAGACCTATTCGCTTTCGTAGCTTGTATGACGGCTG |
| $\mathrm{Arm}_{3,\mathrm{chol}}$ | GTGTTGAGTAGTGAGATGTTCAGCCGTCATACAAGCTACGTTTCGCGAGTGCGGAGTCGTTGC |
| $\mathrm{Arm}_{3,\mathrm{toc}}$ | GGTTTGTTGTTGTGTTGGTTCAGCCGTCATACAAGCTACGTTTCGCGAGTGCGGAGTCGTTGC |
| $\mathrm{Arm}_{\mathrm{block,D_b}}$ | CCAACACAACAACAAACC |
| $\mathrm{Arm}_{\mathrm{block,B_b}}$ | CATCTCACTACTCAACAC |
| A1 | GGTTTGTTGTTGTGT/FluorT/GGAAACTGACAGACCTATTCGC |
| A1* | GCGAATAGGTCTGTCAGTTT |
| A1*-A2 | GCAACGACTCCGCACTCGCGGCGAATAGGTCTGTCAGTTT |
| A2* | CGCGAGTGCGGAGTCGTTGC |
| A2*-A3 | CGCGAGTGCGGAGTCGTTGCCGTAGCTTGTATGACGGCTG |
| A3* | CAGCCGTCATACAAGCTACG |

### References

(1) Angelova, M. I.; Dimitrov, D. S. Liposome electroformation. *Faraday Discuss. Chem. Soc.* **1986**, *81*, 303–311.

(2) Angelova, M. I.; Soléau, S.; Méléard, P.; Faucon, F.; Bothorel, P. Preparation of giant vesicles by external AC electric fields. Kinetics and applications BT - Trends in Colloid and Interface Science VI. Darmstadt, 1992; pp 127–131.

(3) Veatch, S. L.; Keller, S. L. Separation of Liquid Phases in Giant Vesicles of Ternary Mixtures of Phospholipids and Cholesterol. *Biophysical Journal* **2003**, *85*, 3074–3083.

(4) Zadeh, J. N.; Steenberg, C. D.; Bois, J. S.; Wolfe, B. R.; Pierce, M. B.; Khan, A. R.; Dirks, R. M.; Pierce, N. A. NUPACK: Analysis and design of nucleic acid systems. *Journal of Computational Chemistry* **2010**, *32*, 170–173.

(5) Kaufhold, W. T.; Brady, R. A.; Tuffnell, J. M.; Cicuta, P.; Di Michele, L. Membrane Scaffolds Enhance the Responsiveness and Stability of DNA-Based Sensing Circuits. *Bioconjugate Chemistry* **2019**, *30*, 1850–1859.

(6) Stellwagen, N. C.; Magnusdottir, S.; Gelfi, C.; Righetti, P. G. Measuring the translational diffusion coefficients of small DNA molecules by capillary electrophoresis. *Biopolymers* **2001**, *58*, 390–397.

(7) Yurke, B.; Mills, A. P. Using DNA to Power Nanostructures. *Genetic Programming and Evolvable Machines* **2003**, *4*, 111–122.

(8) Zhang, D. Y.; Winfree, E. Control of DNA Strand Displacement Kinetics Using Toehold Exchange. *Journal of the American Chemical Society* **2009**, *131*, 17303–17314.

(9) Czogalla, A.; Petrov, E. P.; Kauert, D. J.; Uzunova, V.; Zhang, Y.; Seidel, R.; Schwille, P. Switchable domain partitioning and diffusion of DNA origami rods on membranes. *Faraday Discussions* **2013**, *161*, 31–43.

(10) Khmelinskaia, A.; Mücksch, J.; Petrov, E. P.; Franquelim, H. G.; Schwille, P. Control of Membrane Binding and Diffusion of Cholesteryl-Modified DNA Origami Nanostructures by DNA Spacers. *Langmuir* **2018**, *34*, 14921–14931.
